# Sparse epistatic patterns in the evolution of terpene synthases

**DOI:** 10.1101/822544

**Authors:** Aditya Ballal, Caroline Laurendon, Melissa Salmon, Maria Vardakou, Jitender Cheema, Paul E. O’Maille, Alexandre V. Morozov

## Abstract

We explore sequence determinants of enzyme activity and specificity in a major enzyme family of terpene synthases. Most enzymes in this family catalyze reactions that produce cyclic terpenes – complex hydrocarbons widely used by plants and insects in diverse biological processes such as defense, communication, and symbiosis. To analyze the molecular mechanisms of emergence of terpene cyclization, we have carried out in-depth examination of mutational space around (E)-β-farnesene synthase, an *Artemisia annua* enzyme which catalyzes production of a linear hydrocarbon chain. Each mutant enzyme in our synthetic libraries was characterized biochemically, and the resulting reaction rate data was used as input to the Michaelis-Menten model of enzyme kinetics, in which free energies were represented as sums of one-amino-acid contributions and two-amino-acid couplings. Our model predicts measured reaction rates with high accuracy and yields free energy landscapes characterized by relatively few coupling terms. As a result, the Michaelis-Menten free energy landscapes have simple, interpretable structure and exhibit little epistasis. We have also developed biophysical fitness models based on the assumption that highly fit enzymes have evolved to maximize the output of correct products, such as cyclic products or a specific product of interest, while minimizing the output of byproducts. This approach results in a non-linear fitness landscape which is considerably more epistatic. Overall, our experimental and computational framework provides focused characterization of evolutionary emergence of novel enzymatic functions in the context of micro-evolutionary exploration of sequence space around naturally occurring enzymes.

## Introduction

Quantitative understanding of the molecular mechanisms of protein evolution is a major challenge in evolutionary biology and protein engineering. The availability and the diversity of evolutionary paths leading to proteins with novel biochemical functions are ultimately determined by a complex pattern of energetic interactions between amino acid residues within the protein, as well as protein-ligand interactions [4-7]. The emergence of novel catalytic functions is a paramount example of evolutionary expansion with profound biological implications. Here we focus on the evolution of ring-forming reactions in terpene synthases (TPSs), a major family of enzymes found in a variety of plants and insects [9, 10]. Cyclic terpenes comprise hundreds of stereochemically complex mono- and polycyclic hydrocarbons; they are involved in pollination, plant and insect predator defense mechanisms, and symbiotic relations. They are also widely used as flavors, fragrances and medicines; a well-known example of the latter is artemisinin, a naturally occurring anti-malarial drug extracted from *Artemisia annua*. Terpenes and terpenoids are the primary constituents of many essential oils in medicinal plants and flowers; examples include α-bisabolol, a monocyclic sesquiterpene alcohol which forms the basis of a colorless viscous oil from German chamomile (*Matricaria recutita*) and *Myoporum crassifolium*, and zingiberene, a monocyclic sesquiterpene that is the predominant constituent of ginger oil.

Enzymes in the TPS family are capable of converting several universal substrates into a diverse variety of terpene products. For example, amorpha-4,11-diene synthase (ADS) produces amorpha-4,11-diene, the bicyclic hydrocarbon precursor of artemisinin, from farnesyl pyrophosphate (FPP), a linear substrate. From an evolutionary point of view, the emergence of the terpene cyclization mechanism served as a crucial step toward creating a major family of enzymes capable of producing diverse and complex metabolic products. Thus, understanding of the evolutionary mechanisms and pathways leading to novel TPS products will enable us to gain deeper insights into enzyme evolution and molecular evolution in general.

In order to investigate TPS evolution in *A. annua* systematically, we have previously used structure-based combinatorial protein engineering (SCOPE) [11] to construct a library with ADS mutations within 6 Å of the active site of (E)-β-farnesene synthase (BFS), which catalyzes the conversion of FPP into the linear hydrocarbon (E)-β-farnesene, an aphid alarm pheromone. BFS shares 49% amino acid sequence identity with ADS [1]. A subset of mutants in the library was characterized biochemically in terms of its spectrum of product terpenes (using gas chromatography – mass spectrometry, GC-MS) [12, 13] and the total kinetic rate of substrate conversion into product (using the malachite green assay, MGA) [14]. Although we have not observed significant amorpha-4,11-diene production in any of the mutants, we have identified several TPSs which produce sizable quantities of α-bisabolol – a major product of TPS enzymes in *A. annua* and *Asteracea* plants. Therefore, in this work we have focused on a highly specific α-bisabolol-producing mutant which contains 5 mutations with respect to the *A. annua* BFS. Specifically, we used SCOPE to create a library of 2^5^ proteins containing all combinations of mutations that bridge BFS and the novel α-bisabolol producing enzyme, BOS, thereby constructing a complete map of all mutational pathways that connect the two enzymes in the subspace in which each of the 5 amino acids can be either in the wild-type or mutant state. Through our mutagenesis studies, we discovered a single-residue substitution (T429A) that imparts a robust and specific α-bisabolol synthase activity in the presence of the T402L gateway mutation which we previously identified as a key mutation necessary to activate cyclization [1].

To characterize the biophysical landscape of our mutant libraries, we have developed a novel approach using the Michaelis-Menten model of enzyme kinetics [15, 16], which has allowed us to express enzymatic reaction rates in terms of the corresponding free energies. Similar to spin-glass models widely used in statistical mechanics [17, 18], we have expanded the free energies in terms of one- and two-body (pairwise) energetic parameters, which correspond to single-amino-acid contributions and amino-acid couplings, respectively. We have fit the model to the available enzyme kinetic rate data and demonstrated that it is capable of both reproducing experimental measurements of kinetic rates with high accuracy and making novel predictions. Thus, there is no need to include higher-order terms such as three-body interactions. Moreover, the number of non-zero pairwise terms is low, making free energy landscapes surprisingly easy to model and interpret. We have used our spin-glass-like models of Michaelis-Menten landscapes to develop a hierarchy of biophysical models of enzyme fitness, interpreting the latter in terms of the protein’s ability to catalyze reactions beneficial to the cell while minimizing production of deleterious or unwanted by-products. We have found that, compared to the free energy landscapes, biophysical fitness landscapes are more epistatic. Nonetheless, the extent of epistasis and landscape roughness are relatively limited in our synthetic libraries, which do not explore the full spectrum of aa substitutions and focus on mutations in the immediate vicinity of the naturally occurring *A. annua* BFS sequence. Overall, we find that the emergence of terpene cyclization can be explained using compact, interpretable models with a relatively small number of free parameters.

## Results

### Elucidation of residue substitutions essential for α-bisabolol synthesis

By sampling natural sequence variation in the background of *A. annua* (*E*)-β-farnesene synthase (BFS), we elucidated residue networks underlying the emergence of terpene cyclization in *A. annua* [1]. Through the biochemical characterization of numerous enzyme variants in this effort, we consistently observed cyclic terpene products present among the broader *Asteraceae* TPS enzyme family. Chief among products with detectable levels of production in several enzyme variants was α-bisabolol, a cyclic terpene alcohol. One variant that contained five amino acid substitutions produced especially high levels of α-bisabolol (61%) (Fig. 1A). This observation was notable, given that α-bisabolol is the product of a dedicated TPS enzyme in *A. annua* and in other *Asteracea* plants. *A. annua* BFS is most closely related to α-bisabolol synthase (BOS) from *Matricaria recutita* and the two enzymes share 70% sequence identity at the amino acid level, consistent with their close evolutionary relationship.

**Figure 1.**
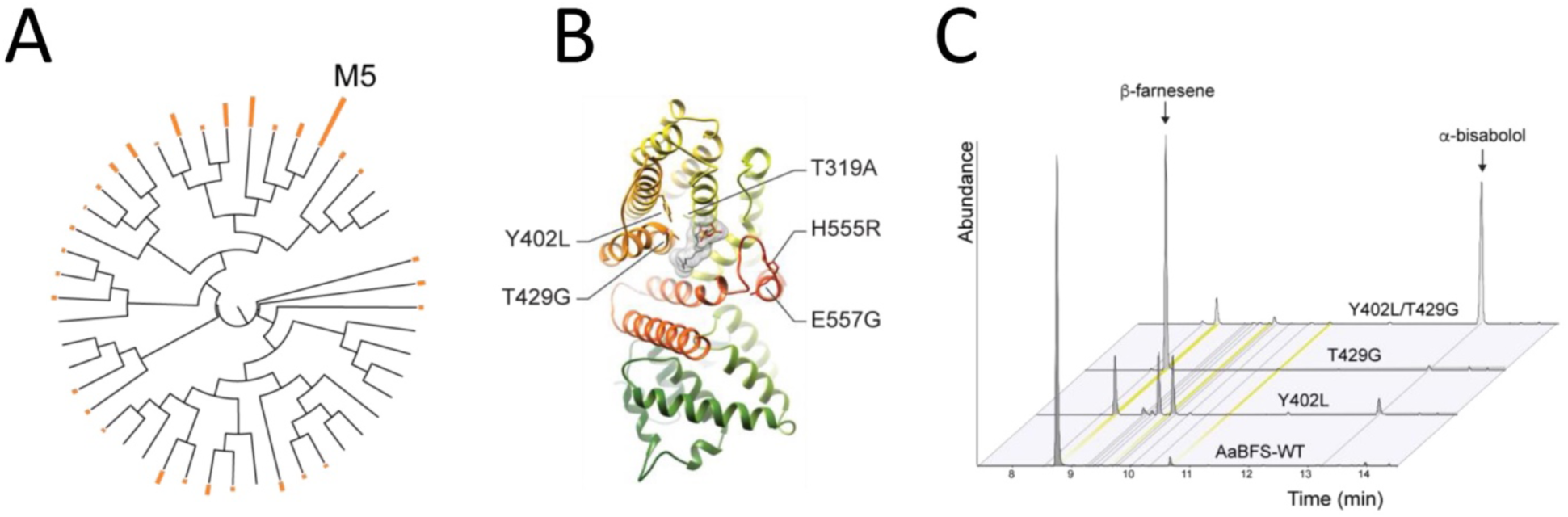
Discovery of a-bisabolol synthase activity in the *A. annua* BFS library. (A) A phylogenetic tree of BFS variants from Salmon et al. [1] was created using ClustalW [53]. The resulting tree was annotated using the Interactive Tree of Life [54] according to the percentage of α-bisabolol products produced (orange bars). The M5 mutant is labeled. (B) Structural positions of residue substitutions in the M5 mutant used for the M5 library synthesis and characterization. (C) GC chromatograms of select members of characterized mutants, with major products indicated.

To identify which residues were responsible for α-bisabolol activity, we designed a library consisting of all combinations of the 5 amino acid mutations (2^5^ = 32 sequences in total) bridging BFS and our previously discovered α-bisabolol-producing BFS variant (M5) (Fig. 1A,B). Using SCOPE, we synthesized the M5 library and verified clones by sequencing. Next, we characterized recombinant enzymes for product specificity by GC-MS [12, 13] and measured their kinetic properties using MGA [14]. Consistent with the mutants from the *A. annua* BFS 6 Å library [1], the Y402L substitution was essential for product cyclization: in the absence of Y402L, all mutants produced linear terpene products. Of the cyclic-producing variants, two product profiles were evident, either a multiple product profile (as seen with the Y402L single mutant) or α-bisabolol as the dominant product in the profile (Fig. 1C). We found that α-bisabolol product specificity was primarily attributable to a single additional mutation T429G in the Y402L background (Fig. 1C), whereas the presence of additional mutations (T319A, H555R and E557G) had weaker effects on product specificity. However, kinetic analysis revealed that the total kinetic rate *k*_*cat*_ was affected significantly by the additional mutations in the T429G/Y402L background. In particular, the C-terminal H555R mutation was very detrimental to catalytic activity. Alone, H555R resulted in a 66% reduction in enzyme activity compared to the BFS wild-type (BFS-WT) enzyme; and in combination with the other mutations, enzyme activity was further reduced to between 1 and 6% of BFS-WT activity. In comparison, the α-bisabolol-producing Y402L/T429G mutant (69% of the total output is α-bisabolol) has moderate catalytic activity (26% of BFS-WT activity) comparable to other native and specific TPS enzymes. Guided by these results, we sought to build quantitative models of enzyme kinetics and energetics, utilizing both the M5 library and the larger collection of BFS mutants from our original study [1].

### Quantitative description of enzyme libraries

We consider two libraries of mutant enzymes in this work. Each enzyme in the library can have mutations at up to *L* variable positions compared to the wild-type sequence, such that the amino acid alphabet at each position is allowed to be in one of the two states: wild type (*W*) or mutant (*M*). Thus, each enzyme sequence *S*_*j*_ can be represented by

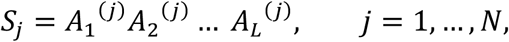

where *A*_*k*_^(*j*)^ = [*W, M*] is the amino acid at position *k* in sequence *j* and *N* is the total number of sequences in a given library. Note that *k* numbers variable positions rather than absolute amino acid positions within a sequence, and that the invariant part of the sequence outside of the *L* positions is not considered explicitly. For each enzyme in both libraries, reaction rates for *n* = 11 distinct products (*k*_*cat,i*_, *i* = 1, …, *n*) have been measured (several other products had negligibly low rates and are therefore excluded from this study). The first library, which we shall refer to as the M25 library, contains *k*_*cat,i*_ measurements for 93 distinct sequences, including the wild-type, with mutations at up to 25 variable positions [1]. The second library, which was described above as the M5 library, contains 2^5^ = 32 sequences, including the wild-type, for all possible combinations of mutations at 5 positions (319, 402, 429, 555, 557) which separate BFS from the novel α-bisabolol producing enzyme found in the M25 library. The combined library contains *N* = 122 distinct sequences, including BFS-WT, with mutations at ≤ 25 variable positions.

### Enzyme kinetics

We have modeled enzymatic reaction rates using the Michaelis-Menten model of enzyme kinetics ([15, 16]). According to this model, enzymes catalyze chemical reactions in a two-step process. The first step is a reversible reaction where a substrate molecule binds the enzyme’s active site. In the second reaction, assumed to be irreversible, substrate is transformed into product and released from the enzyme. In general, terpene synthases in our libraries catalyze multiple reactions simultaneously starting from the same substrate. We assume that the first step is the same in all these reactions since it involves just the substrate and the enzyme:

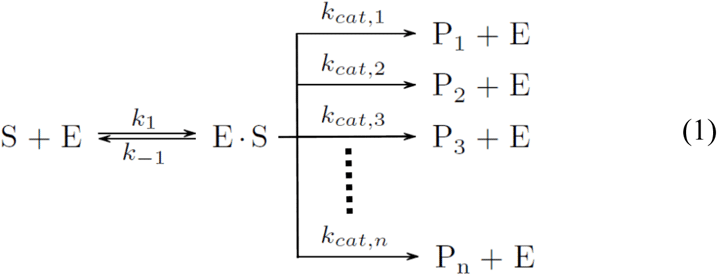

Here, S is the substrate, E is the enzyme, P_i_’s are the products, and *k*_*cat,i*_ denotes the reaction rate for product *i*. Each reaction rate *k*_*cat,i*_ of an enzyme with sequence *S*_*j*_ depends on the Gibbs free energies *G*_3_ and *G*_4,*i*_ (Fig. 2A):

**Figure 2.**
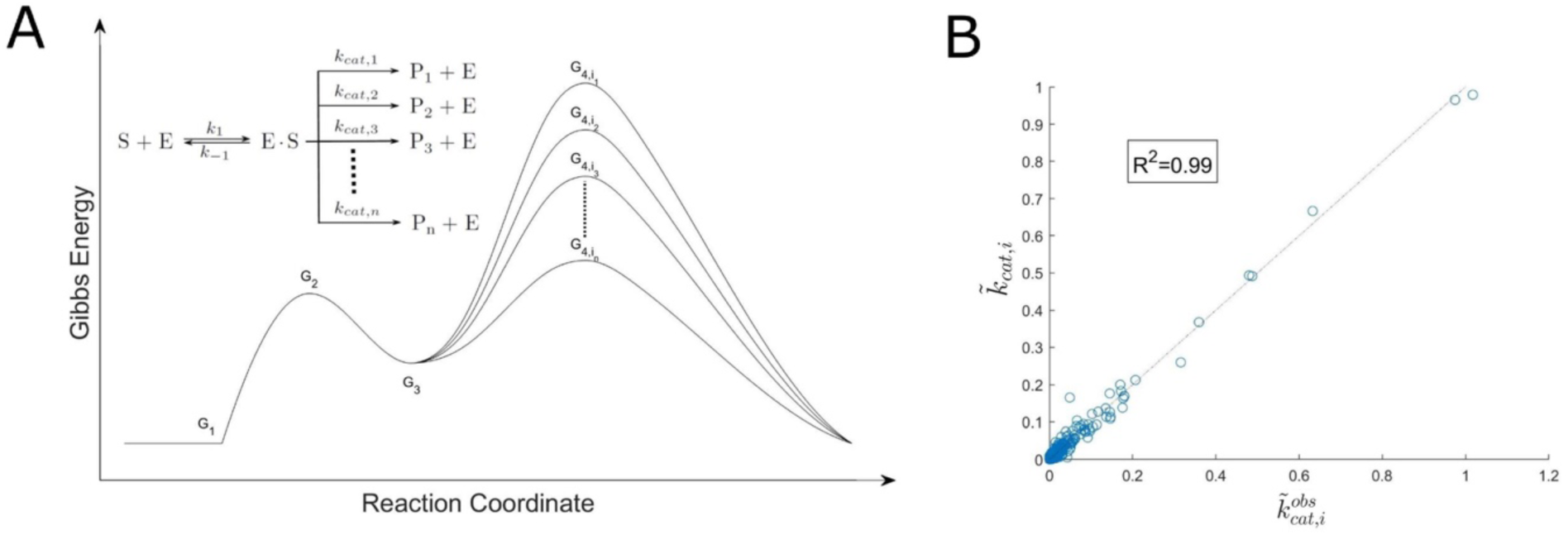
Reaction rate prediction with the pairwise model. (A) Michaelis-Menten model of enzyme kinetics. Shown are free energy profiles for converting substrate S into products *P*_1_ … *P*_*n*_, catalyzed by the enzyme). *G*_1_, *G*_2_, *G*_3_ and 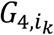 are free energies at the various stages of the enzymatic reactions, and 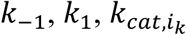 are the corresponding reaction rates as shown in the inset (product indices *i*_1_ … *i*_*n*_ are sorted in the decreasing order of 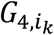 in the panel). Each reaction rate 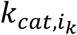 depends on the difference between free energies *G*_4,*i*_ and *G*_3_ (Eq. (2)). Inset shows the corresponding kinetic rates of the Michaelis-Menten reaction. (B) Michaelis-Menten reaction rates predicted using the pairwise model with cross-validation (see Materials and Methods for details).

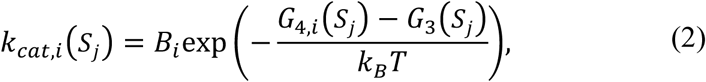

where *B*_*i*_ is the reaction rate for product *i* in the absence of the free energy barrier, *k*_*B*_ is the Boltzmann constant, and *T* is the temperature. As discussed above, we assume that *G*_3_ is independent of the product for a given enzyme, while *G*_4,*i*_ is product-specific. Note that the total reaction rate is given by *k*_*cat*_(*S*_*j*_) = ∑_*i*_ *k*_*cat,i*_ (*S*_*j*_) and that the probability *c*_*i*_ of producing product *i* is therefore *c*_*i*_(*S*_*j*_) = *k*_*cat,i*_(*S*_*j*_)/*k*_*cat*_(*S*_*j*_). Thus, the observed reaction rates are given by 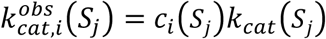 where *c*_*i*_(*S*_*j*_) can be inferred from GC-MS data and *k*_*cat*_(*S*_*j*_) is measured using MGA (Table S1).

### Inference of enzyme energetics with a pairwise model

To model enzyme kinetics and energetics, we have employed a pairwise model inspired by spin-glass models in statistical physics [17, 18]. Such models have been used extensively to study protein stability and protein-protein interactions [19-22]. Unlike these previous approaches, which typically use protein sequence alignments as input to generating novel sequences and scoring the existing ones, our model is designed to predict reaction rates *k*_*cat,i*_ as a function of the enzyme’s sequence. For a given enzyme, we represent the Gibbs free energies *G*_3_ and *G*_4,*i*_ for each product *i* as the sum over single-amino-acid (aa) terms and two-amino-acid coupling terms:

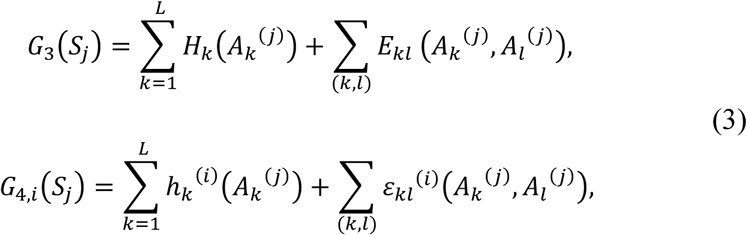

where *H*_*k*_/*h*_*k*_^(*i*)^ and *E*_*kl*_/*ε*_*kl*_(^*i*^) are the one-aa and two-aa contributions to *G*_3_ and *G*_4,*i*_, respectively, and the (*k, l*) sum in the second term on the right-hand side runs over all pairs of variable amino acids in each sequence. This is a set of *N*(*n* + 1) = 1464 equations for the combined dataset. To reduce the number of parameters, we set all terms containing one or two wild-type amino acids to zero. This guarantees that all *G*_3_ and *G*_4,*i*_ values are zero automatically for wild-type sequences, while leaving enough degrees of freedom to model one-aa effects (through the *H*_*k*_(*M*)/*h*_*k*_^(*i*)^(*M*) terms) and two-aa couplings (through the *E*_*kl*_(*M, M*)/*ε*_*kl*_^(*i*)^(*M, M*) terms). For the combined library and for all Gibbs free energies *G*_3_ and *G*_4,*i*_, this procedure yields up to (*n* + 1) = 12 non-zero one-aa terms at each of *L* = 25 positions and, similarly, up to 12 non-zero two-aa coupling terms at each of *L*_*p*_ = 138 pairs of positions. To avoid overfitting, we fit the model using a modified version of the MATLAB implementation of the LASSO method [23, 24] with separate penalties for one-body terms and two-body couplings. The optimal values of penalty term prefactors were obtained using 4-fold cross validation (see Materials and Methods for details).

We find that our fitting procedure yields sparse solutions for Michaelis-Menten free energy landscapes. Indeed, our collection of models for *G*_3_ and *G*_4,*i*_ fitted to the combined data set with optimal one-body and two-body penalties contains a total of 113 out of (*n* + 1)*L*=300 possible one-body terms and just 54 out of (*n* + 1)*L*_*p*_ = 1656 possible two-body couplings, that is, on average, 9.4 out of 25 possible one-body terms and 4.5 out of 138 possible two-body couplings per free energy landscape (see Table S2 for all model parameters). Thus, the observed *k*_*cat,i*_ values of 11 products for 122 different sequences (1342 *k*_*cat,i*_ values in total) are described using just 178 parameters: 113 one-body terms, 54 two-body couplings, and 11 sequence-independent offsets *C*_*i*_ (see Materials and Methods for details). The model fits the reaction rate data with *R*^2^ = 0.99 (Fig. 2B). Note that we typically report reaction rates relative to 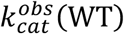, the observed total reaction rate of the wild-type sequence: 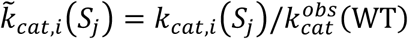 and 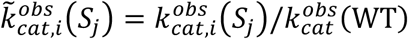 are the predicted and observed relative reaction rates for product *i*.

We observe that at 24 out of 25 positions under consideration (position 559 being the sole exception), mutating a residue results in a change in one or, more typically, several single-aa contributions to free energy landscapes (Fig. S1). Interestingly, changes in *G*_3_ are predominantly negative, meaning that the mutations tend to have adverse effects on the overall values of reaction rates (Eq. (2)). The only exception to this rule is position 402. A Y402L mutation at this position does not just increase the total reaction rate due to the product-independent lowering of the free energy barrier, it also rebalances the enzyme specificity towards the cyclic products, by lowering the relative reaction rates for linear products (E)-β-farnesene (the original BFS product) and nerolidol and increasing the relative reaction rates for cyclic products zingiberene, β-bisabolene, and α-bisabolol (Fig. S1). Thus 402 plays a role of a “gateway” cyclization-unlocking mutation between enzymes producing linear and cyclic products. Other key positions which promote production of cyclic products are 324 and 429. Finally, note that mutations at 10 out of 25 positions result in single-aa terms that suppress (E)-β-farnesene production.

In addition to single-aa terms, the structure of the free energy landscapes is shaped by 54 non-zero coupling terms between pairs of aa positions, 48 of which affect *G*_4,*i*_ values and the other 6 correspond to *G*_3_ (Fig. S2A). Interestingly, most of the *G*_4,*i*_ non-zero couplings contribute to a single free energy landscape corresponding to (E)-β-farnesene – the original linear product of the wild-type BFS enzyme. Out of the 48 two-aa *G*_4,*i*_ terms which determine enzyme specificity, 10 increase reaction rates for cyclic products and only 2 decrease those rates; for linear products, 21 terms contribute to a rate increase and 15 to a rate decrease. Overall, the values tend to be more negative for cyclic products (Fig. S2B), meaning that, as a rule, two-body terms tend to promote cyclization. As shown in Fig. S2C, only 8 aa pairs affect more than one free energy landscape: for example, the 398-429 coupling simultaneously increases the reaction rates of cyclic products α-exo-bergamotene and zingiberene. Correspondingly, 37 aa pairs have a single non-zero coupling term and therefore mutations at these positions affect only one free energy landscape. The remaining 93 pairs of positions do not contribute to the free energies at all.

Since the total number of potentially non-zero fitting parameters is greater than the number of reaction rate measurements, we have carried out additional checks of the validity of the LASSO approach by randomly sampling the parameters of the pairwise model from distributions obtained by fitting the model to the actual *k*_*cat,i*_ data (see Materials and Methods for details). These randomly sampled parameters were subsequently used to generate artificial values *k*′_*cat,i*_ of reaction rates. The input parameters were then recovered using the LASSO approach with cross-validation, identical to the procedure used to fit experimentally observed reaction rates. We were able to accurately predict both the parameters of the model (Fig. S3A) and the corresponding free energy values (Fig. S3B), demonstrating that our approach does not suffer from overfitting.

Finally, we have demonstrated the predictive power of the pairwise model by testing its ability to predict reaction rates of novel enzyme sequences, after training the model on only a part of the available data. Specifically, we have randomly chosen 82 enzyme sequences and trained the model with the LASSO constraint and cross-validation as described above, using *k*_*cat,i*_ values of those sequences as input. The model was subsequently used to predict the *k*_*cat,i*_ values of the remaining 40 enzyme sequences which were not used in training the model, with *R*^2^ = 0.88 (Fig. S4).

### Structure of Michaelis-Menten free energy landscapes

The sparseness of the free energy models described above translates into Michaelis-Menten free energy landscapes with simple and interpretable structure. To illustrate this point, we first focus on the *G*_4_ landscape for the cyclic product α-bisabolol (Fig. 3A-C). In the combined library, this landscape is controlled by 14 one-aa and 3 two-aa model parameters; among one-aa parameters, 5 are above 1 *k*_*B*_*T*, and 3 of those, at positions 324, 402, and 429, enhance relative reaction rates for α-bisabolol by lowering the *G*_4_ barrier (Fig. 3A). In comparison to these leading one-aa contributions, two-aa terms play a secondary role. Consequently, in the M5 library, where the amino acid mutations are restricted to positions 319, 402, 429, 555, and 557, the structure of the free energy landscape is largely determined by the states of amino acids at positions 402 and 429 (a third position, 555, plays a secondary role) (red bars in Fig. 3A). Thus, the *G*_4_ landscape is divided into 4 distinct sectors, with the wild-type BFS sequence (TYTHE at the 5 variable positions) being ≈ 3.2 *k*_*B*_*T* less favorable for α-bisabolol production than the 5-point mutant, ALGRG (Fig. 3B). However, other sequences in the same cluster, such as ALGHG and TLGHG, are characterized by even lower *G*_4_ barriers. In fact, as noted above, it is sufficient to carry out just two mutations, Y402L and T429G, in order to obtain an α-bisabolol producing enzyme.

**Figure 3.**
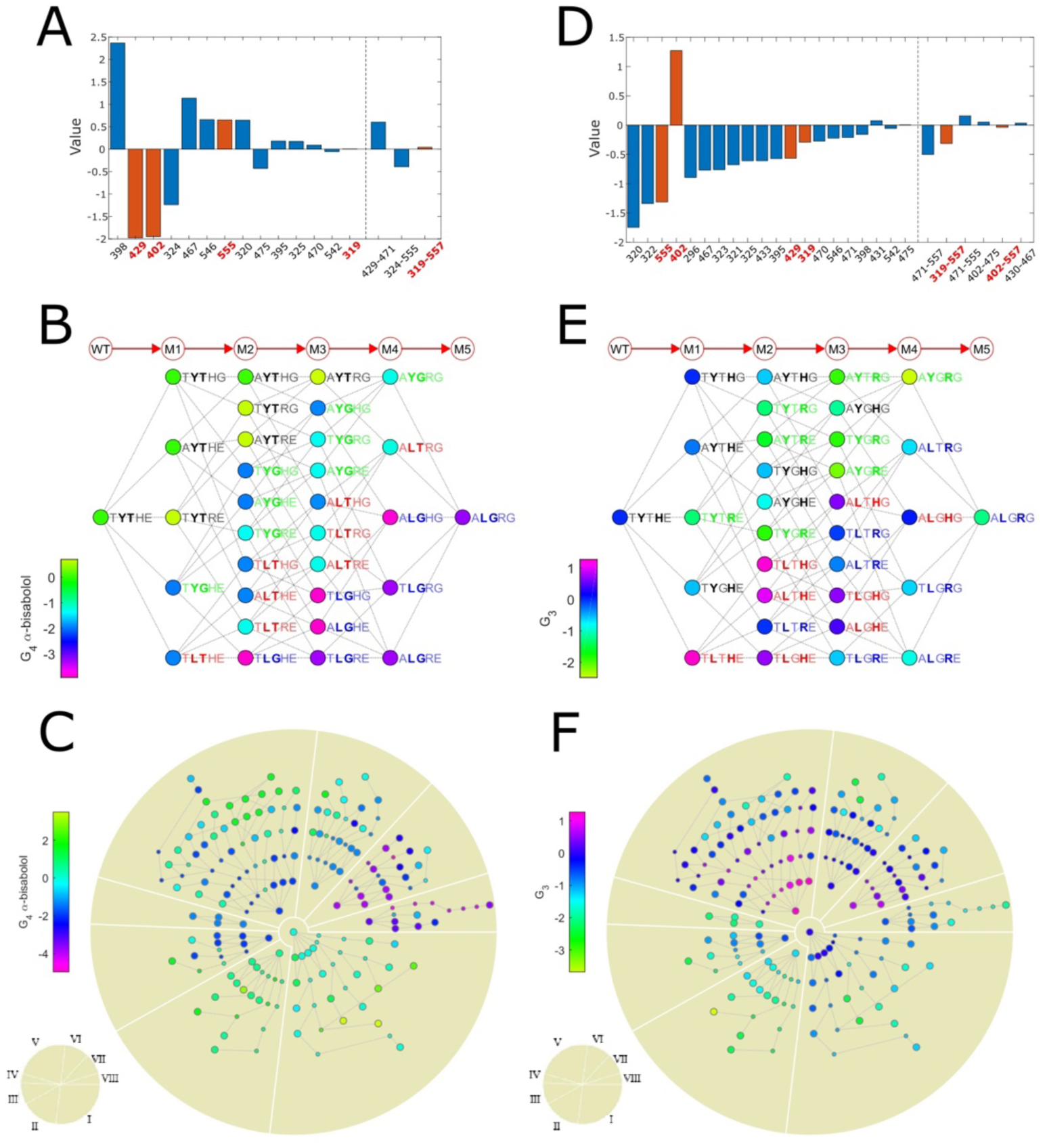
Michaelis-Menten free energy landscapes. (A) The values of all one-aa and two-aa non-zero parameters in the *G*_4_ pairwise expansion for α-bisabolol fitted to the combined library data. Positions and position pairs that occur in the M5 library are highlighted in red. (B) The free energy landscape for α-bisabolol *G*_4_ values on the M5 library, which contains all combinations of mutant and wild-type amino acids at positions 319, 402, 429, 555 and 557. Each node on the landscape is labeled by a string of amino acids at the 5 positions. Nodes that differ by a single amino acid substitution are connected by an edge. The arrows and circles above the landscape indicate the number of mutations away from the wild-type *A. annua* BFS sequence TYTHE. Each node is colored according to the *G*_4_ value for α-bisabolol. In each column, sequences are sorted according to the values of one-aa contributions at positions 402, 429 and 555, which are the 3 largest among the 5 positions considered (Fig. S1). From top to bottom, the sequences with a given number of mutations with respect to the wild-type sequence are sorted in the following order: WWW, WWM, WMW, WMM, MWW, MWM, MMW, MMM. Sequences which differ only at positions 319 and 557 (if any) appear in the order W…W, W…M, M…W, M…M. All nodes are sorted into 4 clusters on the basis of amino acids at positions 402 and 429 which have the largest one-aa contributions (red bars in (A)). These positions are highlighted in bold in each sequence; all sequences are color-coded according to their cluster assignments. (C) Predictions of *G*_4_ for α-bisabolol on the combined library. All nodes are arranged in circles according to the number of mutations away from the wild-type *A. annua* BFS sequence. Nodes are clustered on the basis of positions 402, 429 and 555 in the order WWW, WWM, WMW, WMM, MWW, MWM, MMW, MMM for clusters I-VIII, respectively. These three positions were chosen since they have the highest position score: 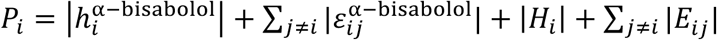, which represents the sum of the absolute magnitudes of all one-aa and two-aa model parameters associated with position *i*. Nodes in the same cluster are connected by an edge if their sequences differ by a single amino acid substitution. Within each cluster and each circular shell, nodes are sorted so as to minimize the number of edges crossing each other. Large circles denote sequences in the combined dataset, whereas small circles show sequences for which *G*_4_ values were predicted. Each node is colored according to the *G*_4_ value for α-bisabolol. (D-F) Same as (A-C) but for the *G*_3_ free energy landscape. Note that in (E), all nodes are sorted as in (B) to enable visual comparisons. However, the nodes are classified into 4 clusters (and their sequences are color-coded) on the basis of amino acids at positions 402 and 555, which have the largest one-aa contributions to *G*_3_ (red bars in (D)). Nodes in (F) are classified into the same clusters and appear in the same order within each cluster as nodes in (C).

Although the specificity of a given enzyme is controlled by the relative heights of the *G*_4_ barriers for each product, its overall output also depends on the height of the *G*_3_ barrier which we have assumed to be independent of the product type. Similar to the *G*_4_ landscape for α-bisabolol, the *G*_3_ free energy landscape in the combined library is a function of just 20 one-aa and 5 two-aa model parameters, with only 4 one-aa parameters, at positions 320, 322, 555, and 402, above 1 *k*_*B*_*T* (Fig. 3D). Two of these positions, 402 and 555, are variable in the M5 library and hence the amino acid states at these positions largely determine the structure of the *G*_3_ free energy landscape (red bars in Fig. 3D and Fig. 3E; positions 429 and 319 play a secondary role). Interestingly, although positions 319 and 557 are characterized by small (319) and zero (557) one-aa contributions, they shape the free energy landscape through a two-aa term. We observe that the 5-point mutant, ALGRG, is lower in the overall output than the wild-type sequence, TYTHE: the corresponding *G*_3_ value is lower by ≈ 1.2 *k*_*B*_*T*, which makes the kinetic barrier higher overall. The Y402L mutation is the only major contributor to lowering the free energy barrier (Fig. 3D); consequently, the TLGHE double mutant discussed above (Y402L/T429G) favors both α-bisabolol production and high overall output.

To investigate the effect of these mutations on other products, we have considered *G*_4_ free energy landscapes for the wild-type linear product, (E)-β-farnesene, and another cyclic product of practical importance, zingiberene (Fig. S5). Interestingly, the (E)-β-farnesene landscape is characterized by 17 one-aa and 28 two-aa terms and thus can be expected to be more epistatic (Fig. S5A), although its projection onto M5 sequences is fairly sparse, being mainly determined by aa states at positions 402 and 429 (Fig. S5B). As expected, the TLGHE double mutant (Y402L/T429G) and especially the 5-point mutant, ALGRG, are characterized by sharply decreased levels of (E)-β-farnesene production. Correspondingly, the best (E)-β-farnesene producing enzymes are those with aa at positions 402 and 429 left in the wild-type state. The *G*_4_ free energy landscape for zingiberene is determined almost exclusively by one-aa contributions (Fig. S5D), of which aa states at positions 402, 429, and 555 structure the landscape’s projection onto M5 sequences. Since the Y402L mutation is the only one favorable for zingiberene production, sequences in the “LT” cluster (shown in red in Fig. S5E), which includes a single mutant TLTHE, are best zingiberene producers.

It is also informative to consider the free energy landscapes on all sequences from the combined library. In Fig. 3C, nodes are arranged radially around the wild-type sequence according to the number of mutations. Sequences are clustered into 8 sectors on the basis of aa states at positions 402, 429 and 555, which contribute the most when both one-aa and two-aa terms are taken into account. In addition to 122 sequences in the combined dataset, we have made predictions for 69 additional sequences which were chosen to fill in the gaps in mutational pathways connecting experimentally characterized sequences. Clusters VII and VIII, which have both Y402L and T429G mutations (cluster VII has H whereas cluster VIII has R at position 555), are enriched the most in α-bisabolol producing enzymes (Fig. 3C). These clusters, along with clusters V and VI, tend to contain sequences that are least favorable for (E)-β-farnesene production due to the Y402L mutation (Fig. S5C). For zingiberene, the most favorable sequences are concentrated in cluster V, although the *G*_4_ free energy barrier is rarely lowered by more than 1 *k*_*B*_*T* (Fig. S5F). Finally, sequences with higher values of *G*_3_ (which is beneficial for the overall output) tend to be found in clusters V and VII (Fig. 3F). In summary, sequences in cluster VII (Fig. 3C,F) are the best candidates for α-bisabolol production in the combined library; specific candidates can be chosen based on how critical it is to produce α-bisabolol specifically (as opposed e.g. to a mixture of α-bisabolol, zingiberene, and other cyclic products).

### Epistasis on free energy landscapes

The notion of epistasis is closely related to the extent of non-linearity and ruggedness observed in fitness or energy landscapes [25-27]. In its most basic form, the idea of epistasis concerns aa states at two distinct positions in the sequence. In the case of two aa states (such as the *W* and *M* states considered in this work), epistatic interactions allow for a simple geometric interpretation (Fig. 4A). Note that the no-epistasis scenario implies the absence of a coupling between the two sites, while the other three scenarios (magnitude, sign or reciprocal sign epistasis) are controlled by the magnitude and the sign of the relevant coupling terms. In our case, only the *MM* coupling can be non-zero by construction; however, the two aa sites in question are embedded into longer sequences and therefore the amount and the type of epistasis are also affected by the two-aa terms in which one of the partners is outside of the current pair.

**Figure 4.**
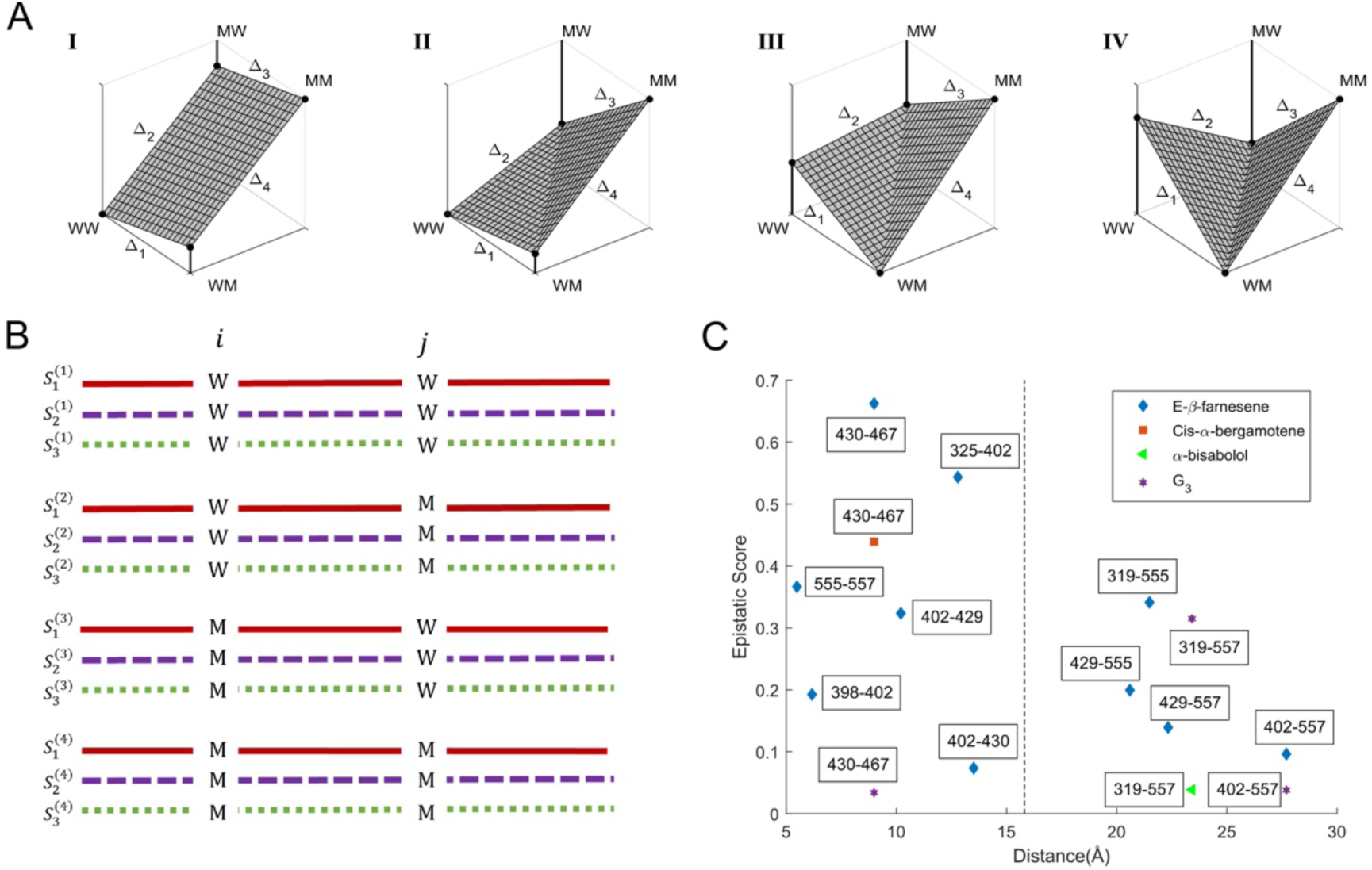
Epistasis on free energy landscapes. (A) The four types of epistasis in a two-site system. The aa at each site can be in either *W* or *M* state. Panel I: no epistasis, with each mutation contributing the same amount to the total free energy (or fitness) regardless of the aa state at the other site. Panel II: magnitude epistasis, where the magnitude (but not the sign) of the aa free energy (or fitness) contribution depends on the aa state at the other site. Panel III: sign epistasis, where for one of the sites, the sign (and, in general, the magnitude) of the aa free energy (or fitness) contribution depends on the aa state at the other site. Panel IV: reciprocal sign epistasis, where the aa free energy (or fitness) contribution at both sites depends on the aa state at the other site. (B) Schematic representation of a set of sequences *S*_*k*_^(*a*)^ divided into 4 subsets *T*_*ij*_(*A, B*) of equal size, where *A, B* = [*W, M*]. The sequence subsets are used to calculate the epistasis score *ES*_*ij*_ for aa positions *i* and *j*, as described in the text. Note that outside of the positions *i* and *j*, sequences in each subset are exactly the same. (C) Plot of the free energy epistasis scores *ES*_*ij*_ vs. the *C*_*α*_ − *C*_*α*_ spatial distances (in Å) between aa positions *i* and *j*. All epistasis scores with magnitudes < 0.01 were excluded from the plot. A vertical dashed grey line at 15.8 Å shows the average *C*_*α*_ − *C*_*α*_ distance for all 138 aa pairs considered in this work.

Since the magnitude and the sign of epistasis between two aa sites depend on the rest of sequence, we have focused our attention on the subset of sequences which are identical outside of two positions *i* and *j*. At these positions, data has to be available for all 4 aa states: (*W, W*), (*W, M*), (*M, W*) and (*M, M*). These requirements result in 4 sequence subsets of the same size: 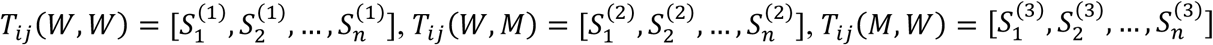, and 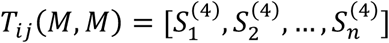 (Fig. 4B). Next, we compute the free energies 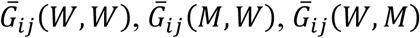, and 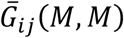 which are simply the *G*_3_ or *G*_4_ free energy values averaged over all sequences in the corresponding *T*_*ij*_ subset. Finally, we define the differences of the averaged free energies along each edge in the geometric shapes of Fig. 4A: 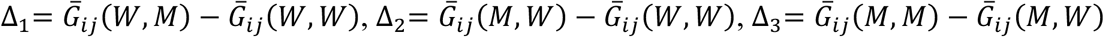, and 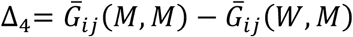. Then the four epistasis types can be succinctly summarized as follows: Δ_1_= Δ_3_ and Δ_2_= Δ_4_ correspond to the absence of epistasis, otherwise sgn(Δ_1_) = sgn(Δ_3_) and sgn(Δ_2_) = sgn(Δ_4_) represent magnitude epistasis, sgn(Δ_1_) ≠ sgn(Δ_3_), sgn(Δ_2_) = sgn(Δ_4_) or sgn(Δ_1_) = sgn(Δ_3_), sgn(Δ_2_) ≠ sgn(Δ_4_) represent sign epistasis (the two cases correspond to two opposite pairs of edges in Fig. 4A, panel III), and sgn(Δ_1_) ≠ sgn(Δ_3_), sgn(Δ_2_) ≠ sgn(Δ_4_) correspond to the reciprocal sign epistasis. Note that on fitness landscapes, sign epistasis can significantly affect genotype accessibility by making some evolutionary trajectories unavailable or unlikely [26], whereas reciprocal sign epistasis is a necessary condition for the existence of multiple local maxima [28].

Next, we define an epistatic score *ES*_*ij*_ as the absolute magnitude of the difference between the Δ values on the two pairs of opposite edges in the geometric shapes shown in Fig. 4A:

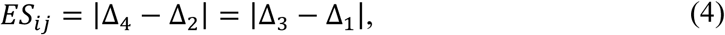

such that *ES*_*ij*_ = 0.0 in the case of no epistasis, and positive otherwise.

Consistent with the above discussion of Michaelis-Menten landscape structure and appearance, all free energy landscapes are characterized by a limited amount of epistasis. Indeed, in our combined dataset we find only 16 pairs of positions *i* and *j* for which the above analysis of epistatic interactions can be carried out: 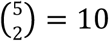 pairs come from the M5 library, with 2^3^ = 8 sequences in each *T*_*ij*_ subset, 1 pair (402-430) has 3 sequences in each *T*_*ij*_ subset, and 5 more pairs occur on the background of a single sequence. Since there are 12 distinct free energy landscapes, we have 192 potentially epistatic instances in our dataset. Out of those, only 15 pairs are characterized by *ES*_*ij*_ *>* 0.01 *k*_*B*_*T*, with 13 exhibiting magnitude epistasis, 1 showing sign epistasis (positions 430-467 on the *G*_3_ landscape, with *ES*_430-467_ = 0.03 *k*_*B*_*T*), and 1 more demonstrating reciprocal sign epistasis (positions 430-467 on the *G*_4_ landscape for cis-α- bergamotene, with *ES*_430-467_ = 0.44 *k*_*B*_*T*) (Table S3). Interestingly, 10 out of 13 pairs with magnitude epistasis occur on the (E)-β-farnesene landscape, and 2 other instances occur on the *G*_3_ landscape. Thus, (E)-β-farnesene production mediated by a wild-type enzyme is characterized by significantly more pronounced epistatic interactions than production of cyclic terpenes by the enzymes in our mutant library.

To investigate whether higher levels of free energy epistasis occur in pairs of residues that are close to each other in 3D space, we have plotted the 15 pairs of residues with non-zero *ES*_*i*)_ scores vs. the corresponding *C*_*α*_ − *C*_*α*_ distances in Fig. 4C. Although the two residues in pairs with top 3 *ES*_*i*)_ values do tend to be closer to each other than 15.8 Å, the average distance between all 138 pairs under consideration (cf. the vertical dashed line in Fig. 4C), the overall trend is rather weak, especially if all pairs that are close to each other in the linear sequence, such as 555-557, are excluded from the consideration. For example, the 319-555 pair on the (E)-β-farnesene landscape exhibits significant magnitude epistasis, despite the fact that these residues are separated by more than 20 Å. Remarkably, the 430-467 pair appears as epistatic on 3 free energy landscapes, with the corresponding *ES*_*i*)_ scores ranked 1, 3, and 15 by absolute magnitude (out of 15 pairs with *ES*_*i*)_ 5 0.01 *k*_*B*_*T*; cf. Table S3).

Strong epistasis between positions 430 and 467 can be readily rationalized by their spatial proximity in the BFS structural model (Fig. 5A). The residues in the 430-467 pair are within van der Waals distance (<3 Å) from each other and are located at the bottom of the active site, making direct contacts with the isopropenyl tail of the substrate FPP. As such, substitutions at these positions are expected to be interdependent, with hydrophobic contacts likely accounting for the physical interactions between the two residues, at least in the wild-type.

**Figure 5.**
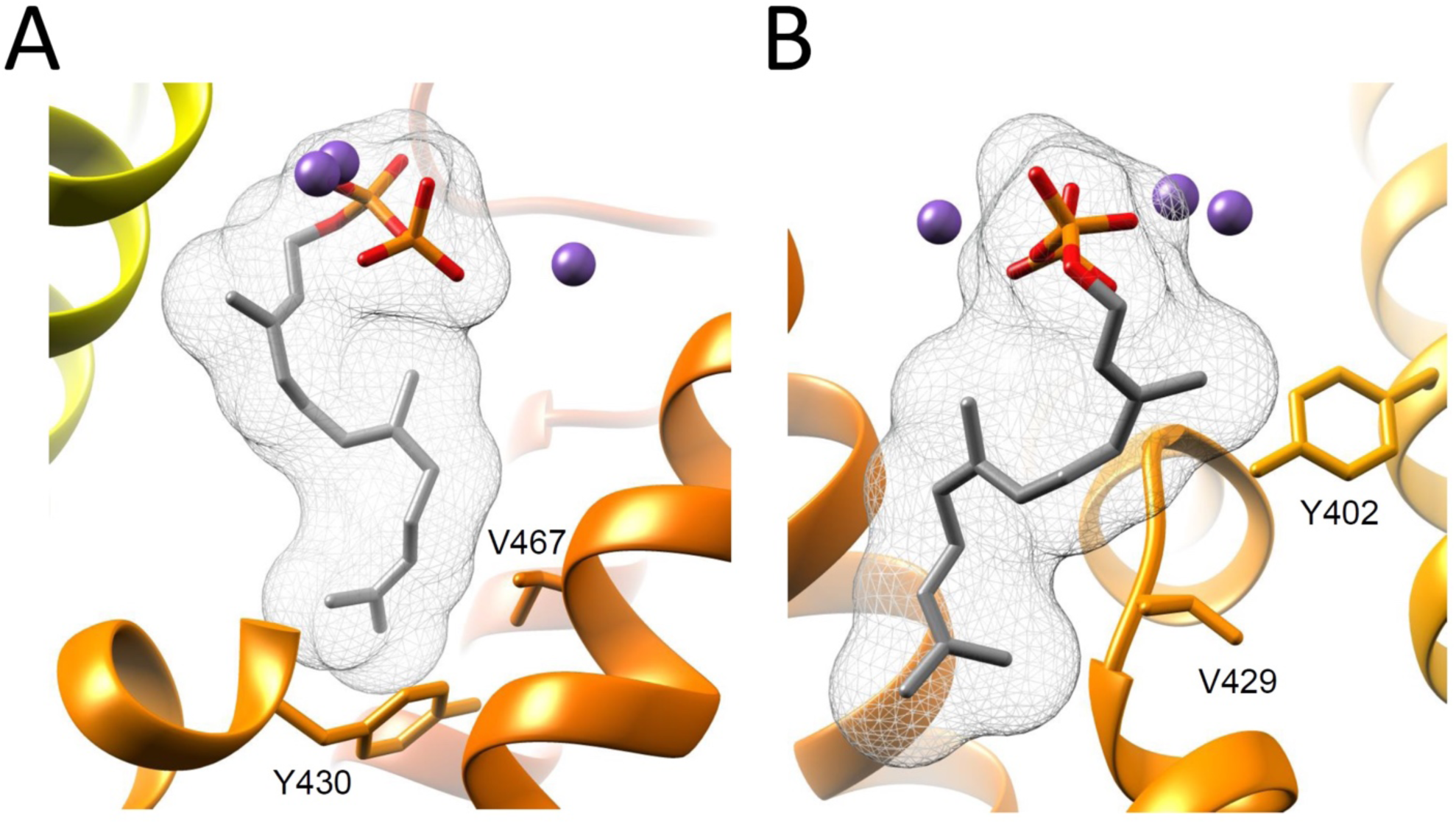
Structural basis of epistasis on free energy landscapes. Shown are ribbon diagram cut-outs of the BFS structural homology model (created in I-TASSER [55, 56]) with docked FPP substrate (mesh) [1]. Magnesium ions (purple spheres) coordinate the pyrophosphate moiety at the top of the active site. (A) Amino acids at positions 430 and 467 interact with each other and with the isopropenyl tail of the substrate at the bottom of the active site. (B) Amino acids at positions 402 and 429 are in spatial proximity with each other and with the isopropenyl tail of the substrate.

### Fitness models

We have developed a class of biophysical fitness landscapes based on the Michaelis-Menten theory of enzyme kinetics and energetics. We start with an assumption that all products produced by a given enzyme can be classified into correct and incorrect. Making a correct product results in a fitness gain while making an incorrect product results in a fitness loss due to the necessity of its removal or degradation. In a biotechnology setting, products are classified into correct and incorrect by the researcher in a context of a specific project, whereas in a cellular setting the needs of the cell and the associated fitness gains and losses may be time-or environment-dependent. Thus, in general a single enzyme’s fitness *F* is given by a weighted sum over fitness gains and losses associated with each product:

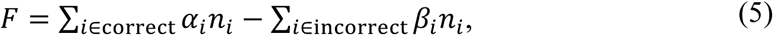

where *n*_*i*_ is the number of product molecules of type *i* produced per unit time and *α*_*i*_, *β*_*i*_ are the corresponding fitness gains and losses per product molecule. Note that 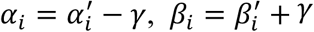, where *γ* > 0 is the fitness cost of making or acquiring a substrate molecule (for example, *γ* is expected to be close to 0 if the substrate molecules are abundant in the environment). We assume that *α*_*i*_ > 0, ∀*i* or, in other words, that benefits of making the correct products outweigh all the associated costs (products that are less valuable than substrates can be accounted for in the second term on the right-hand side). In the absence of information on product-specific rewards and penalties, we set all fitness gains and losses to be product-independent, which makes the fitness a weighted difference between the total number of correct and incorrect product molecules produced per unit time:

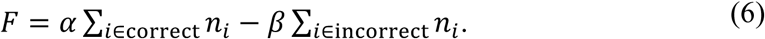

Note that in a cellular setting, fitness gains and losses may depend on the number of produced molecules: *α*_*i*_ = *α*_*i*_(*n*_*i*_), *β*_*i*_ = *β*_*i*_(*n*_*i*_). For example, fitness may be maximized only if a given molecule’s production rate is close to optimal; overproduction may lead to diminished returns and even sign reversal. Although such extensions are easy to model within our framework, here we focus on the product type- and product rate-independent scenario (Eq. (6)). For simplicity, we label all cyclic products as correct and all non-cyclic products as incorrect; alternative scenarios such as a single correct product (favoring specific enzymes typically found in nature) can be easily considered.

Within the Michaelis-Menten framework, *n*_*i*_ is given by the reaction velocity per enzyme molecule:

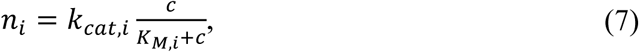

where *K*_*M,i*_ is the Michaelis constant and *c* is the substrate concentration. Thus,

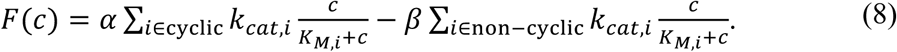

In the high substrate-concentration limit (*c* ≫ *K*_*M,i*_, ∀*i*), the enzyme velocity reaches its maximum value and the expression for fitness becomes

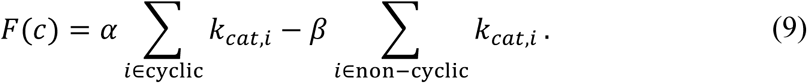

In this limit, fitness is simply a function of the enzyme’s reaction rates and is independent of substrate concentration. In the low substrate-concentration limit (*K*_*M,i*_ ≫ *c*, ∀*i*),

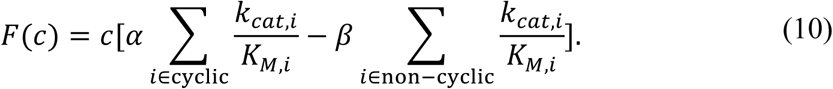

In this case, fitness is proportional to the substrate concentration *c* and depends on both reaction rates and Michaelis constants. Moreover, if the height of all the *G*_3_-*G*_4,*i*_ barriers is low, such that *k*_*cat,i*_ ≫ *k*_-1_, ∀*i*, fitness becomes approximately independent of the reaction rates and the product type: 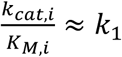, and the overall enzyme velocity is determined by the height of the *G*_1_-*G*_2_ free energy barrier (Fig. 2A).

Note that if we do not know anything about substrate concentration *a priori*, we can assume that it is uniformly distributed in the [*c*_*min*_, *c*_*max*_] range. Then the expected value of *F* is given by

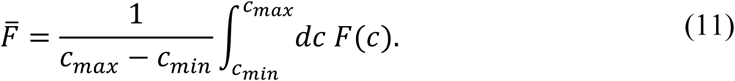

If *c*_*max*_ ≫ *K*_*M,i*_, ∀*i*, the integral in Eq. (11) is dominated by the high substrate-concentration limit and we again recover Eq. (9). Therefore, we focus on the high substrate-concentration case (Eq. (9)) in the subsequent analysis.

### Structure of the fitness landscape and epistatic interactions

We have computed fitness values for each sequence in the combined library using Eq. (9). Since the overall scale of each term and therefore the absolute magnitude of the fitness contribution cannot be determined from our analysis alone, we have set *α* = 1, *β* = 1 and have all shifted fitness values such that the fitness of the wild-type BFS sequence, TYTHE, is exactly zero (see Fig. 6A for the fitness landscape on the M5 library). We have chosen to use predicted rather than observed reaction rates in Eq. (9); switching to the experimentally observed rates would have made little difference since our model predicts reaction rates *k*_*cat,i*_ with high accuracy (Fig. 2B).

**Figure 6.**
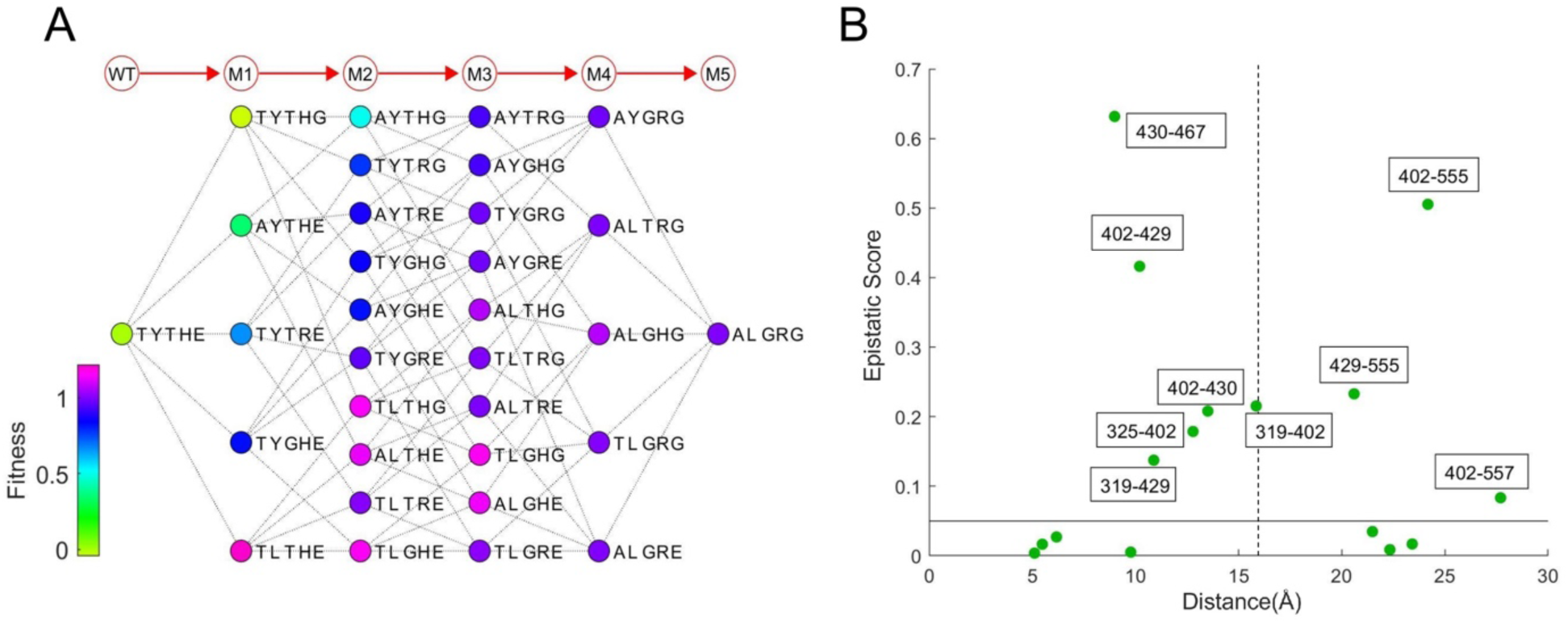
Michaelis-Menten fitness landscape. (A) Fitness landscape for the M5 library. All fitness values were computed using Eq. (9) with *α* = 1, *β* = 1 and predicted (rather than observed) reaction rates *k*_*cat,i*_. The landscape is presented as in Fig. 3B, with the nodes sorted in the same order to facilitate visual comparisons. Each node is colored according to its fitness value. (B) Plot of the fitness epistasis scores *ES*_*ij*_ vs. the *C*_*α*_ − *C*_*α*_ spatial distances (in Å) between aa positions *i* and *j*. A horizontal solid black line indicates the 0.05 cutoff separating residue pairs considered non-epistatic from the rest. A vertical dashed grey line at 15.8 Å shows the average *C*_*α*_ − *C*_*α*_ distance for all 138 aa pairs considered in this work.

Similar to the free energy landscapes discussed above, the fitness landscape exhibits simple structure: creation of cyclic products is enabled by a single gateway mutation, Y402L, so that a single-point mutant, TLTHE, exhibits a significant jump in fitness. In fact, sequences with L at position 402 and G or T at position 429 are the best producers of cyclic products; however, the 5-point mutant, ALGRG, has somewhat lower fitness, largely because its overall reaction rate is lower (Fig. 3E). The relatively simple structure of the fitness landscape is somewhat expected since the reaction rates in Eq. (9) depend on the *G*_3_ and *G*_4_ free energy values, which are determined by just a few non-zero two-aa terms. On the other hand, fitness is a non-linear function of the free energies, which can lead to epistasis even if the underlying free energy model has no two-aa terms at all [29, 30].

To study the amount of epistasis on our fitness landscape, we have computed epistatic scores *ES*_*i*)_ using Eq. (4) for the 16 pairs of positions identified earlier in the epistatic analysis of free energy landscapes (Fig. 6B). The only difference with the previous analysis is that fitness values rather than free energy values were averaged in each *T*_*i*)_ subset. We have categorized all pairs with *ES*_*i*)_ < 0.05 as exhibiting no epistasis. Out of the remaining 9 pairs, 8 show sign epistasis and 1 exhibits reciprocal sign epistasis (Table S3), indicating that the fitness landscape is indeed rougher than the free energy landscapes considered earlier and, as a consequence, single-point mutation evolutionary trajectories are expected to be constrained. We do not observe a prominent correlation between fitness epistatic scores and the corresponding *C*_*α*_ − *C*_*α*_ distances (Fig. 6B): although the 430-467 and 402-429 pairs ranked first and third by the absolute magnitude of the epistatic score are separated by less than the average distance between all aa pairs, the residues in the 402-555 pair, which is ranked second, are nearly 25 Å apart. Strikingly, the 402-555 pair does not contribute to epistasis on any of the free energy landscapes (Fig. 4C, Table S3). Thus, considerable long-range couplings can be created purely through non-linearities in the Michaelis-Menten fitness function. In contrast, the 430-467 pair is the most significant contributor to epistatic interactions on the free energy landscapes and its residues are in direct contact with one another (Fig. 4C, Fig. 5A).

Similar to the 403-467 pair, residues 402 and 429 are within 5 Å of one another and make direct contacts with the FPP substrate (Fig. 5B). Given their shared role in forming one side of the active site, the residues in the 402-429 pair are positioned to influence substrate folding and guide product formation. The role of residue 402 in activating cyclization stems from its interaction with the first isoprene unit of the substrate, which enables an initial isomerization reaction. Residue 429 resides deeper in the binding pocket and therefore more likely influences substrate folding. Together, the 402-429 pair enables cyclization of a substrate conformation that readily undergoes 1,6 cyclization while the reaction is terminated by water capture, likely associated with magnesium ions positioned near the mouth of the active site. In sum, the fact that the 402-429 pair shows strong epistasis in cyclic product formation is entirely consistent with its structural role in the active site.

Given the large distance between residues at positions 402 and 555 (≈ 25 Å), we thought to rationalize the potential physical basis for interactions in the 402-555 pair through a network analysis of the BFS protein structure, whose purpose is to delineate the intervening interactions (Fig. S6). The network analysis identified 3 shortest pathways with 4 edges, likely not mutually exclusive, that connect residues 402 and 555 (Fig. S6B). Pathway 1 involves interactions exclusively between aa in the protein structure, while pathways 2 and 3 depend on intervening connections through the isopropenyl chain of the FPP substrate. Interestingly, pathway 2 transits through 429, the gateway residue that controls cyclic product synthesis. Pathway 3 transits through position 327, a conserved catalytic aspartic acid of the DDxxD motif. Subtle misalignment of the pyrophosphate-magnesium complex coordinated by the DDxxD motif, propagated through this interaction network, may provide an explanation for reduced catalytic efficiency observed upon aa substitution at position 555.

## Discussion and Conclusion

In this work, we presented synthesis and characterization of a library of mutant terpene synthases designed to explore evolutionary mechanisms underlying emergence of terpene cyclization. We explored the mutational space of this major enzyme family using BFS from *A. annua*, which catalyzes production of the linear hydrocarbon (E)-β-farnesene, as a starting point. Each enzyme in the library was characterized biochemically using gas chromatography – mass spectrometry [12, 13] and the malachite green assay [14], enabling us to obtain reaction rates for 11 products, 7 of which are cyclic. The library is a combination of two independent datasets: a previously published partial map of mutational pathways connecting BFS to ADS, which catalyzes production of amorpha-4,11-diene, a bicyclic molecule [1], and a complete map of mutational pathways between BFS and BOS, an α-bisabolol-producing synthetic enzyme identified in the BFS-to-ADS screen. The combined library has 122 enzyme sequences, with amino acids mutated at ≤ 25 positions compared to BFS. At each variable position, the corresponding aa can only be either in the wild-type (BFS) or mutant state, resulting in the effective aa alphabet of size 2. In-depth biochemical characterization of this enzyme library has enabled us to carry out quantitative modeling of enzyme energetics and evolution.

We have described each enzyme in our library using the Michaelis-Menten model of enzyme kinetics [15, 16]. The Michaelis-Menten framework allows us express enzymatic reaction rates and the overall reaction velocity in terms of free energies assigned to various enzymatic states (Fig. 2A). These free energies are closely related to the free energies of protein folding and binding which have been extensively explored using protein engineering methods [31-33], with ΔΔ*G* data available for multiple proteins [34]. These studies reveal that effects of multiple mutations on protein energetics are nearly additive, especially if the mutations are distant from each other in the linear sequence [35]. Consequently, the assumption of independent energetic contributions of residues at different sites has been extensively used in biophysical models of protein evolution that express organismal fitness in terms of protein energetics [30, 36-39]. In the light of these previous findings, we expected Michaelis-Menten free energies to be approximately additive as well, with two-aa coupling terms playing a secondary role. To check this hypothesis, we have represented the free energies as a sum of one- and two-aa contributions which we treated as fitting parameters. The resulting model, supplemented with a LASSO constraint which is designed to minimize the number of non-zero fitting parameters [24], was fit to the reaction rate data, reproducing it with a high degree of accuracy (Fig. 2B).

As expected, the model has very few non-zero two-aa terms: on average, just 4.5 out of 138 coupling parameters contribute to a given free energy landscape. For example, the α-bisabolol *G*_4_ landscape is controlled by 3 coupling parameters (Fig. 3A). Interestingly, the (E)-β-farnesene *G*_4_ landscape is by far the most non-additive as it is characterized by 32 non-zero coupling terms, followed by the *G*_3_ landscape with 6 couplings (Fig. S2A). Since both of these landscapes affect reaction rates of the linear product of BFS, a wild-type enzyme, it is conceivable that the corresponding free energies have been evolutionarily fine-tuned, resulting in a more coupled, non-linear landscape. This implies that novel enzymatic functions can evolve quickly through pathways that do not require establishment of intricate networks of aa interactions right away. However, these networks start to play an increasingly prominent role as the novel enzyme is further optimized by evolution for efficiency and specificity.

We have sought to quantify the extent of ruggedness on the Michaelis-Menten free energy landscapes by considering epistatic interactions between various aa pairs. The notion of epistasis, including higher-order epistasis, has been extensively studied in the context of fitness landscapes [20, 40-42]. Epistasis can profoundly alter evolutionary dynamics on fitness landscapes by restricting the availability of evolutionary trajectories [26] and by creating local maxima that can trap or slow down evolving populations [28]. We have extended the idea of epistasis to the free energy landscapes; as with fitness, aa pairs have been classified into no-epistasis, magnitude epistasis, sign epistasis, and reciprocal sign epistasis categories (Fig. 4A) [27]. We have also introduced an epistatic score *ES*_*ij*_ which is zero in the case of no epistasis and positive otherwise (Eq. (4)). Note that in general this score depends not only on the magnitude of the two-aa term between residues *i* and *j*, but also on the coupling to the residues outside of the *i, j* pair. Consistent with the previous observations in the literature described above and the simple, easily interpretable structure of the free energy landscapes constructed in this work (Fig. 3, Fig. S5), we find free-energy epistasis to be of relatively limited importance, with only 15 out of 192 pairs considered exhibiting any epistasis at all, 13 of which in the magnitude epistasis category. These findings appear to be at variance with a recent report in which significant epistasis was observed in antibody-antigen binding free energies [43]. However, we note that our analysis is effectively limited to the 2-letter alphabet and therefore our observations may change once the full spectrum of mutations is included. Furthermore, enzyme evolution, and specialized metabolic enzymes in particular, may be subject to different constraints than evolution in the adaptive immune system. Finally, our analysis is likely affected by the fact that, by design, we explore sequence space around a naturally occurring enzyme, BFS. Thus, our findings reflect “micro” rather than “macro” evolution which can be studied e.g. on the basis of protein sequence alignments involving multiple protein families and multiple organisms. Analysis of such alignments tends to yield much less interpretable models characterized by numerous non-zero coupling terms [20, 44-49]. Interestingly, although there is a certain degree of enrichment for spatial proximity in strongly epistatic pairs, many of such pairs are separated by >15 Å (Fig. 4C), conceivably as a result of long-range allosteric interactions mediated by networks of intervening amino acids.

Even if the underlying free energy model is purely additive, the corresponding biophysical fitness function may be characterized by epistasis and local maxima if it is a non-linear function of the free energies, as observed in models that include protein folding stability and binding affinity as explicit fitness determinants [30, 37]. We have significantly extended this prior work by constructing a biophysical fitness landscape in terms of free energies of the Michaelis-Menten model. Each enzyme’s fitness is assumed to be proportional to its total reaction velocity for the ‘correct’ product(s) as dictated by a given biological or biotechnological context. Production of incorrect products is penalized in a similar way. Thus, highly tuned enzymes that produce the maximum number of correct molecules and the minimum number of incorrect molecules per unit time would be characterized by high fitness values. Extensions to fitness functions in which e.g. the optimal rates of production of correct products are enforced are straightforward but are not the main focus here.

As expected, the fitness landscape is more epistatic than the free energy landscapes due to its non-linear nature, with 9 out of 16 aa pairs under consideration characterized by significant epistatic scores (8 of these are in the sign epistasis and 1 in the reciprocal sign epistasis category). Thus, the fitness landscape is rougher than its underlying free energy components, which, depending on the balance between selection, mutation, and genetic drift in evolving populations, may render some of the evolutionary pathways inaccessible. Nonetheless, projecting the fitness function onto the M5 library results in a global maximum, TLTHE, and no local maxima (Fig. 6A). Weak correlation between epistatic scores and spatial distances (Fig. 6B) is less surprising in this case since fitness values may depend on the states of residues that are not necessarily energetically coupled, due to compensatory mutations, as for example occurs in biophysical fitness models that depend on the total free energy of protein folding and binding [30, 37, 50].

In conclusion, we have carried out detailed biochemical characterization in a library of mutant enzymes which explore sequence space in the vicinity of BFS, a naturally occurring catalyzer of the linear product (E)-β-farnesene. This characterization has allowed us to construct Michaelis-Menten free energy landscapes, study their structure quantitatively, and use them as input to a simple biophysical model of enzyme fitness. Our analysis highlights a fundamental evolutionary mechanism of creating epistatic interactions through non-linearities in the fitness function and underscores the surprising simplicity and interpretability of enzyme energetics. In the future, we intend to investigate the universality of our findings by employing additional synthetic libraries (in particular, going beyond the 2-letter alphabet explored in this work), and by exploring sequence space around wild-type enzymes from other protein families. Another intriguing direction of future work will be to construct synthetic enzyme libraries that iteratively cover more and more sequence space starting from a wild-type enzyme such as BFS, or attempt to bridge sequence separation between two wild-type enzymes.

## Materials and Methods

### Gene library synthesis

The BFS M5 library (2^5^ = 32 mutants) was constructed in a 3-phase process using structure-based combinatorial protein engineering (SCOPE) which involves (i) fragment amplification, (ii) library recombination, and (iii) library amplification [11]. <u>Fragment amplification.</u> N-terminal and C-terminal gene fragments were amplified by PCR using mutagenized N- and C-terminal plasmid libraries, respectively, as a template. Specific recombination primers and generic amplification primers were designed as described in Dokarry et al. [11]. PCR was carried out using Phusion High-Fidelity polymerase (NEB) using the following protocol: 98 °C for 3 min, followed by 30 cycles of 98 °C for 15 sec, 50 °C for 30 sec, 72 °C for 1 min, and 72 °C for 10 min, then followed by incubation at 4 °C. An aliquot of 2 μl of each reaction was analyzed by agarose gel electrophoresis on a 2% TAE agarose gel, and fragments were diluted 1:10 for use in the SCOPE recombination reaction. <u>Library recombination.</u> Purified, diluted N-terminal and C-terminal fragments were mixed together in a 1:1 ratio and recombined with 1 nM of the central fragment. The reaction was set up as described in Dokarry et al. [11]. Recombination PCR was carried out using Phusion High-Fidelity polymerase (NEB) using the following protocol: 98 °C for 3 min, followed by 30 cycles of 98 °C for 15 sec, followed by a 50-70 °C ramp (50 °C at cycle 1, then +1.5 °C/cycle) for 30 sec, followed by 72 °C for 30 sec, then followed by incubation at 4 °C. Following the reactions, the tubes were stored on ice and used directly in the SCOPE amplification reaction. <u>Library amplification.</u> An aliquot of 2.5 μl of the recombination reaction was used as a template for the SCOPE amplification reaction. The reaction was set up as described in Dokarry et al. [11]. Amplification PCR was carried out using Phusion High-Fidelity polymerase (NEB) using the following protocol: 98 °C for 3 min, followed by 30 cycles of 98 °C for 15 sec, 65 °C for 15 sec, 72 *°C* for 1 min, and 72 °C for 10 min, then followed by incubation at 4 °C. An aliquot of 2 μl of the amplification reaction was analyzed by agarose gel electrophoresis on a 1% TAE agarose gel. Reaction products were PEG-precipitated into the same volume of Tris-EDTA buffer, pH 8.0 (TE buffer) prior to Gateway cloning.

### Cloning of individual mutants

Gateway cloning was carried out in 5 μl reactions. For these reactions, pDONR207 and pH9GW were used as the entry vector and the destination vector, respectively. 1 μl of the BP or LR reaction was transformed into 10 μl of *E. coli* DH5α Library Efficiency cells (Life Technologies) by heat-shock. The transformed cells were spread on LB plates containing antibiotics and incubated at 37 °C overnight. BP clones were confirmed by sequencing to identify mutants of interest prior to the LR reaction. For protein expression, pH9GW plasmids were transformed into 5 μl *E. coli* BL21(DE3) cells (NEB) by heat shock. Following cell recovery in 100 μl Super Optimal Broth (SOC), 10 μl volume of transformed cells was spread on LB plates containing antibiotics and incubated at 37 °C overnight.

### Protein expression of mutants

Single colonies were transferred to 1 ml of liquid media (LB with kanamycin) in 96-well plates, followed by growth for 16 hrs with shaking at 37 °C and 230 rpm. Cultures were diluted 10-fold into 5 ml of Terrific Broth (TB) growth media with kanamycin in 24-well round-bottom plates covered with micro-porous tape, followed by growth with shaking at 37 °C and 180 rpm until cultures reached OD_600_ ≥ 0.8. Protein expression was induced by addition of IPTG to concentration of 0.1 mM, followed by growth with shaking at 20 °C and 180 rpm for 5 hrs. Cells were then harvested by centrifugation and cell pellets were frozen at −20 °C.

### Ni-NTA affinity chromatography purification of library proteins

Pellets from 5 ml expression cultures were re-suspended by adding 0.8 ml of lysis buffer (50 mM Tris-HCl, 500 mM NaCl, 20 mM imidazole, 10% glycerol (v/v), 10 mM β-mercaptoethanol, and 1% (v/v) Tween-20, pH 8) containing 1 mg/ml lysozyme and 1 mM EDTA directly to frozen pellets, followed by shaking at room temperature (25 °C) at 250 rpm for 30 minutes. Subsequently, 10 μl of the benzonase solution (850 mM MgCl_2_ and 3.78 U/μl benzonase (Novagen)) were added, followed by additional shaking at 250 rpm for 15 min. A 400 μl aliquot of lysate was passed through a Whatman unifilter 96-well plate and collected in another Whatman plate containing 50 μl bed-volume (100 μl of slurry) of superflow Ni-NTA resin (QIAgen), pre-equilibrated with lysis buffer using a vacuum manifold. This step was repeated to pass the entire lysate volume through the column. Each well was washed with 1.5 ml lysis buffer (3 × 500 μl), followed by 1.5 ml wash buffer (lysis buffer lacking Tween-20). Resin was air-dried prior to addition of 150 μl elution buffer (wash buffer containing 250 mM imidazole), followed by centrifugation at 1,500 rpm for 2 min to recover eluted protein. The eluate was reapplied to the column and the centrifugation step was repeated. To calculate protein concentrations, 200 μl of Bradford reagent (Bio-Rad) were added to a 10 μl aliquot of purified protein in a flat-bottom 96-well microplate. The reaction was incubated at room temperature for 5 minutes and the OD_595_ was measured on a Varioskan Flash plate reader (Thermo Scientific). Protein concentrations were quantified against a bovine serum albumin (BSA) standard curve.

### Enzyme vial assay

The vial assay was performed in 2 ml screw-top glass vials (Agilent) in a 500 μl reaction volume. Each reaction consisted of assay buffer at pH 7.0 [25 mM 2-(N-morpholino) ethanesulfonic acid (MES), 25 mM N-cyclohexyl-3-aminopropanesulfonic acid (CAPS), 50 mM Tris(hydroxymethyl)aminomethane (Tris)], 5 mM MgCl_2_, 100 μM farnesyl diphosphate (FPP) and enzyme (1.5-3 μM). Reactions were mixed at room temperature and overlaid with 500 μl of hexane (Sigma), and the caps were affixed. After an overnight incubation at room temperature (25 °C), the hydrocarbon products were extracted by vigorous vortexing for 10 sec, followed by GC-MS analysis [12], as described below.

### Product identification and quantification by GC-MS

Reaction products were analyzed using a Hewlett–Packard 6890 gas chromatograph (GC) coupled to a 5973 mass selective detector (MSD) outfitted with a 7683B series injector and autosampler and equipped with an HP-5MS capillary column (0.25 mm i.d. x 30 m with 0.25 μm film) (Agilent Technologies). The sampling depth was set to 7.5 mm, placing the needle in the center of the organic layer (near the 750 μl level in the 2 ml glass vial). The GC was operated at a He flow rate of 0.8 ml/min, and the MSD was operated at 70 eV. Splitless injections of 2 μl were performed with an injector temperature of 250 °C. The GC was programmed with an initial oven temperature of 80 °C (1 min hold), which was then increased at a rate of 20 °C/min up to 140 °C (1 min hold), followed by a 5 °C/min ramp until 170 °C (2 min hold), followed by a 100 °C/min ramp until 300 °C (1 min hold). A solvent delay of 6 min was allowed prior to the acquisition of MS data. Product peaks were quantified by the integration of peak areas using Enhanced Chemstation (version E.02.00, Agilent Technologies). Products were identified using Massfinder 4.25, a 2-D algorithm that employs retention index and mass-spectral fingerprints for compound identification.

The GC-MS data was visually inspected to identify the peaks (compounds) to be quantified in the series of samples. The quantification was carried out automatically and was based on using the mass spectra to obtain chromatograms extracted for ions (m/z) (usually 3-5) specific to each compound. First the intensities of each extracted chromatogram were calculated using Met-Idea v2.05 [51], based on a collection of (RT, m/z) pairs (Table S4). The remainder of the steps were carried out in Matlab 2013 (The MathWorks) using scripts written in-house. For each extracted chromatogram, the intensities were corrected to take into account the percentage signal that the ion represented in the mass spectrum, so that, in a perfect case, the corrected intensities would be the same for all ions and would represent the amount of the compound present (relative quantitation). These intensities were then averaged across ions. The percentage of the signal represented by each compound was then calculated and the outcome saved in a spreadsheet. In addition, a report, from scripts written in-house, was generated which provided a number of useful diagnostic tools, notably graphs showing the extracted chromatograms over the relevant RT range, as well as the correlation between the corrected intensities from different ions. These were used to detect systematic bias resulting from non-specificity and/or interference between closely eluting compounds. Whenever necessary, the list of ions was refined so as to limit such occurrences.

### Malachite green assay for *k*_*cat*_ apparent and steady-state kinetic measurements

Kinetic assays were performed in 96 well flat-bottomed plates (Grenier). For *k*_*cat*_ apparent measurements, assays of 50 μl were conducted in the malachite green assay buffer (vial assay buffer containing 2.5 mU of the coupling enzyme inorganic pyrophosphatase from *S. cerevisiae* (Sigma)) using six 2-fold serial dilutions of the purified protein. Monophosphate (Pi) and pyrophosphate (PPi) standard curves (100 μM to 0.01 μM) were generated using a 2-fold serial dilution in the malachite green assay buffer without FPP. Reactions were set up in duplicate and incubated at room temperature for 30 min, 90 min and 180 min. Enzyme reactions were quenched by addition of the malachite green development solution (prepared according to Pegan et al. [52]), incubated for 15 min, and read at 623 nm on a Varioskan Flash plate reader. For steady-state kinetic measurements, assays of 50 μL were conducted in the malachite green assay buffer using serial dilutions of FPP, with a starting concentration of 100 μM. Enzyme was added to give a final concentration of 0.014 μM, unless otherwise stated. Standard curves of monophosphate (Pi) and pyrophosphate (PPi) (50 μM to 0.01 μM) were generated using serial 2-fold dilutions in the malachite green assay buffer without FPP. Reactions were set up on ice in triplicate and incubated at 30 °C for 15 or 40 min, depending on the mutant studied. Enzyme reactions were quenched by addition of 12 μL of the malachite green development solution, incubated for 15 min, and read at 623 nm on a Varioskan Flash plate reader.

### Second-order model of reaction rates and Michaelis-Menten free energies

Using Eqs. (2) and (3), we obtain a set of *Nn* = 1342 equations for predicting relative reaction rates:

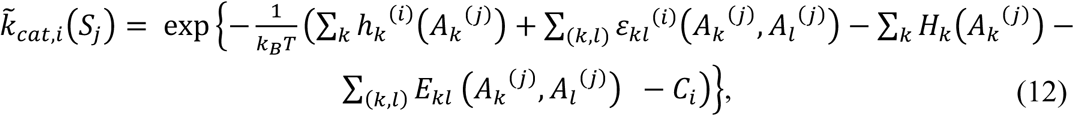

where 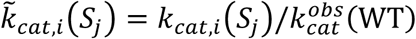 is the predicted relative reaction rate for product *i* 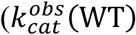 is the observed total reaction rate of the wild-type sequence), and the sequence-independent offsets *C*_*i*_ are defined 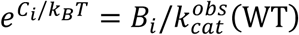 (*B*_*i*_ is the reaction rate for product *i* of the wild-type sequence, cf. Eq. (2)). This set of equations has (*n* + 1)(*L* + *L*_*p*_) + *n* = 1967 fitting parameters since there are *L* one-body terms, *L*_*p*_ coupling terms for each *G*_4,*i*_ and *G*_3_, and one additional term per product for the sequence-independent offset *C*_*i*_. The predicted relative reaction rates are fitted to 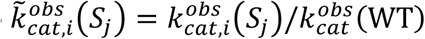, the relative reaction rates observed for each enzyme in the combined library.

Since the total number of fitting variables is greater than the number of measurements, we employ the LASSO (Least Absolute Shrinkage and Selection Operator) algorithm [23, 24], which reduces the number of non-zero fitting parameters by imposing a penalty proportional to their *L*, norm. We impose different penalties on one-body terms and two-body couplings, while the *C*_*i*_ offsets are left unconstrained. The problem therefore reduces to finding a set of fitting parameters which minimize the following expression:

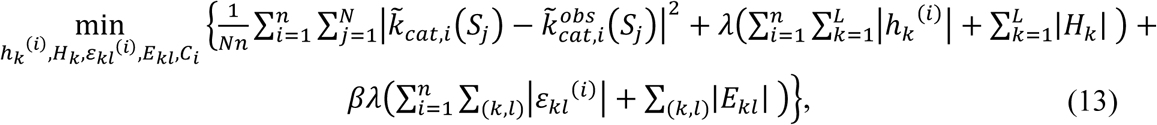

where *λ* and *βλ* are the regularization coefficients which determine the relative importance of the *L*, penalty terms. We have determined the penalty parameters *λ* and *β* by 4-fold cross validation. All enzyme sequences *S*_)_ were randomly partitioned into 4 equal-sized samples. One sample was assigned as the test set and the other 3 as the training set on which the model was fitted. This procedure was repeated 4 times, with each sample used exactly once as the test set. We varied *λ* from 10^−2.8^ to 10^−1.4^ and *β* from 1 to 10. For each pair of *λ* and *β*, we calculated the mean-square error in predicting the test set (the first term in Eq. (13)), and averaged it over all 4 cross-validation runs (Fig. S7). The error was minimized for *λ* = 0.0079 and *β* = 2.7384; these values were subsequently used to fit the model on the entire data set.

### Generation of synthetic data

Since the total number of potentially non-zero fitting parameters is larger than the number of reaction rate measurements in the combined library, we have additionally checked the consistency of the LASSO procedure by fitting our model to artificially generated data. To generate the artificial data, we randomly chose 7 non-zero one-body terms and 4 non-zero couplings for each *G*_4,*i*_ and *G*_3_ landscape and for each enzyme sequence (all other terms were assumed to be zero). These parameters were assigned random values based on two Gaussian distributions which were obtained by computing the mean *m* and the standard deviation σ of the one-body terms (*m* = 0.24, σ = 0.85) and the couplings (*m* = −0.09, σ = 0.63) inferred from the combined library. The sequence-independent offsets *C*_*i*_ were likewise sampled from a Gaussian distribution with *m* = −5.52 and σ = 1.96, where the Gaussian was fit to the set of *C*_*i*_’s obtained after fitting the model to the reaction rate data in the combined library. We have generated artificial 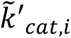 values using these randomly chosen parameters:

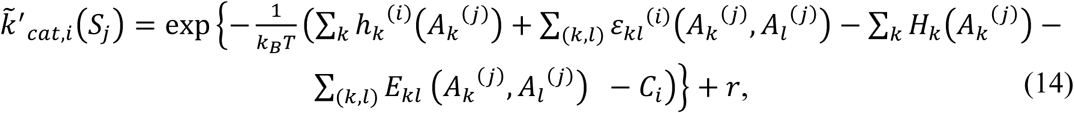

where *r* was randomly sampled from a Gaussian distribution with *m*_*ϵ*_ = 0.0 and σ_*ϵ*_ = 0.003 (σ_*ϵ*_ is the mean-square error obtained after fitting the model to the reaction rate data in the combined dataset). This artificial data set was treated as reaction rate measurements, and the parameters of the model as well as free energy values were subsequently inferred using LASSO as described above (Fig. S3).

## Acknowledgements

PEO acknowledges support from the Biotechnology and Biological Sciences Research Council grant BB/K003690/1 and the Institute Strategic Program Grants BB/J004561/1 (Understanding and Exploiting Plant and Microbial Secondary Metabolism) at JIC and BB/I015345/1 (Food and Health) at IFR. PEO also wishes to thank the John Innes Foundation and the John Innes Centre, Norwich. AVM and PEO acknowledge support from the National Science Foundation (awards MCB1920914 and MCB1920922, respectively).

## Supplementary Figure captions

**Figure S1.**
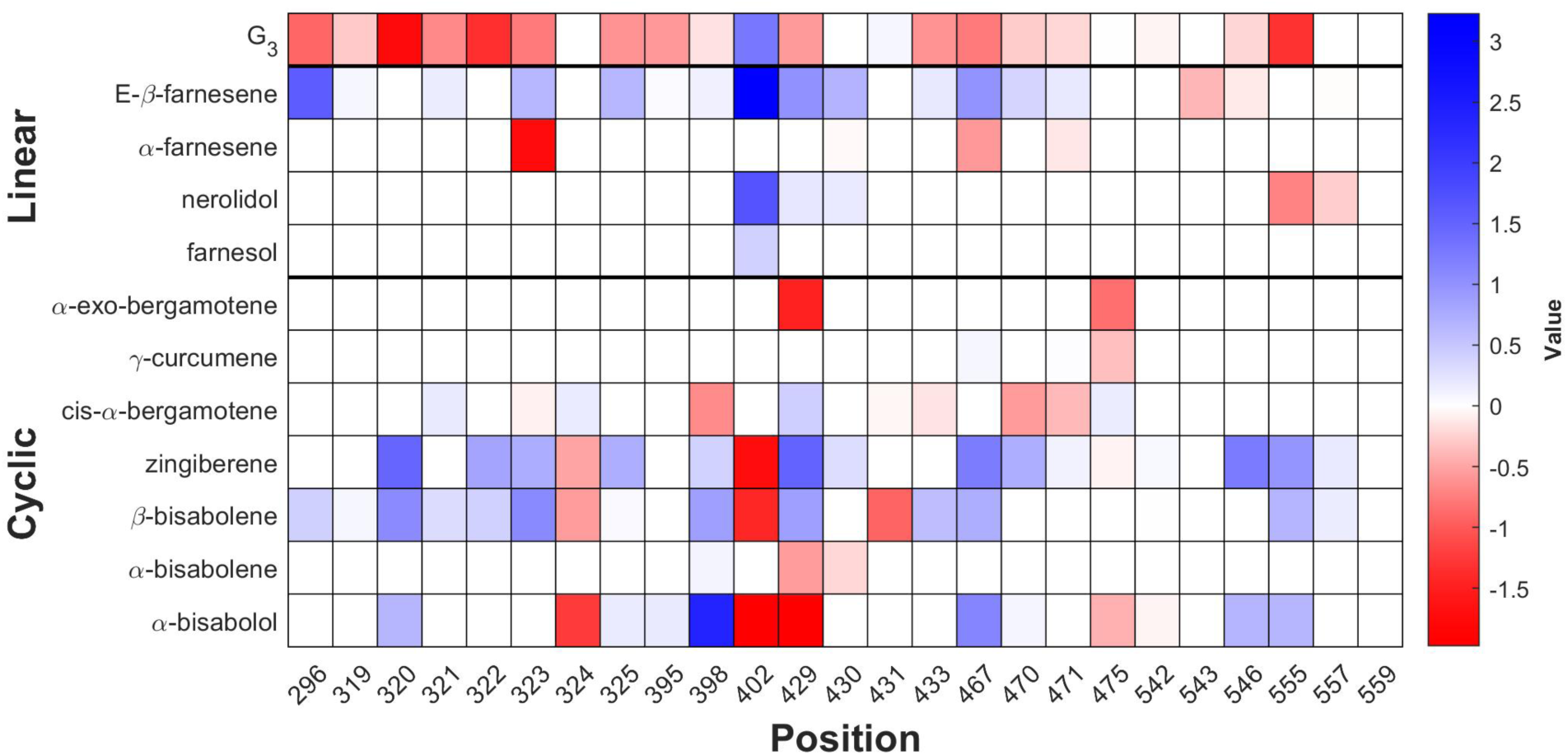
One-aa contributions to Michaelis-Menten free energies. Fitted values of one-aa model parameters *H*_*k*_(*M*) (*G*_3_) and *h*_*k*_^(*i*)^(*M*) (*G*_4,*i*_).

**Figure S2.**
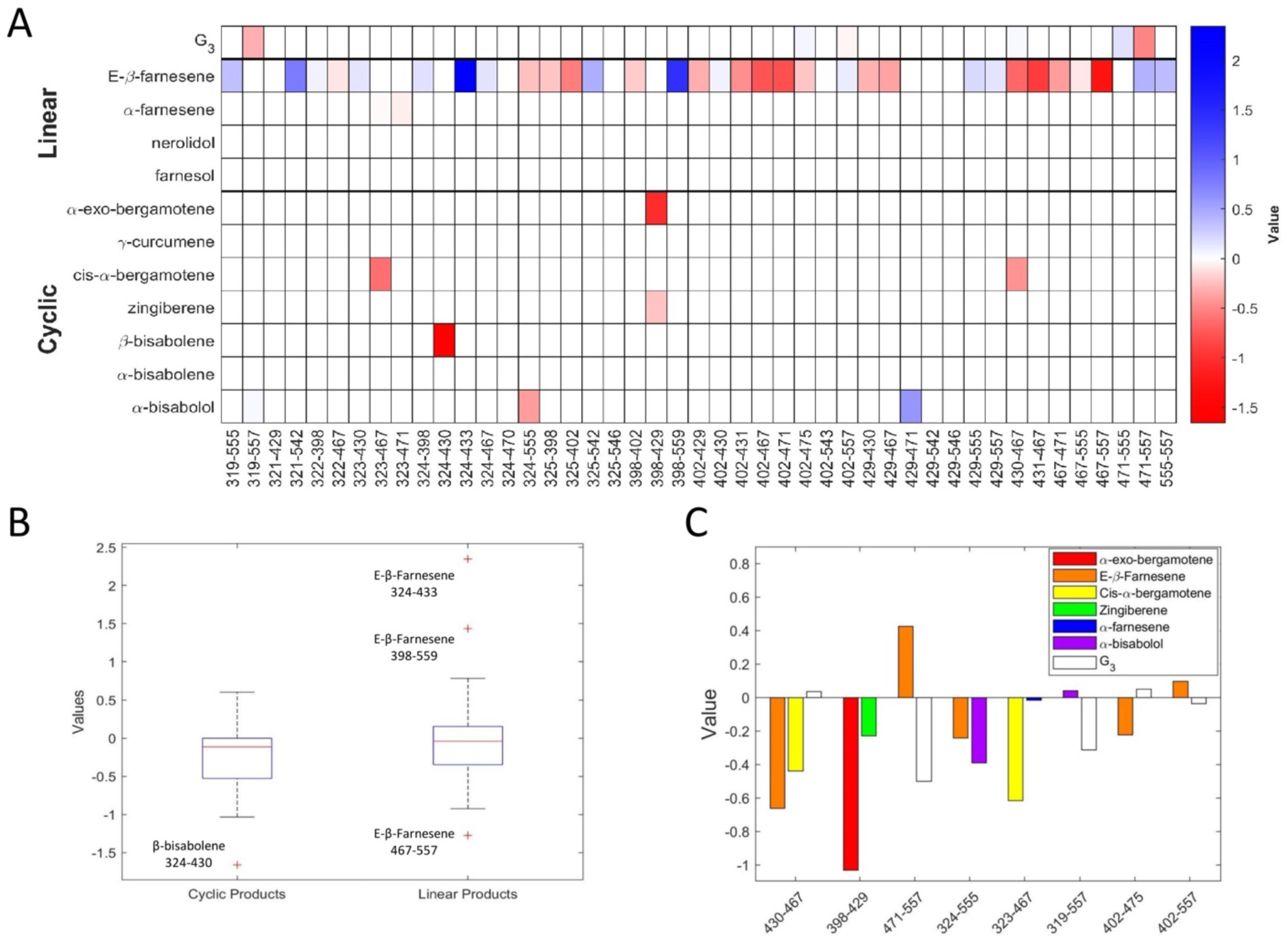
Two-aa contributions to Michaelis-Menten free energies. (A) Fitted values of two-aa model parameters *E*_*kl*_(*M, M*) (*G*_3_) and *ε*_*kl*_^(*i*)^(*M, M*) (*G*_4,*i*_). (B) Box-and-whiskers plots (created using the boxplot function with default parameters in MATLAB) for the distributions of two-aa model parameters for linear and cyclic enzyme products. (C) Two-aa model parameters *E*_*kl*_(*M, M*) (*G*_3_) and *ε*_*kl*_^(*i*)^(*M, M*) (*G*_4,*i*_) at 8 pairs of positions where two or more two-aa contributions adopt non-zero values.

**Figure S3.**
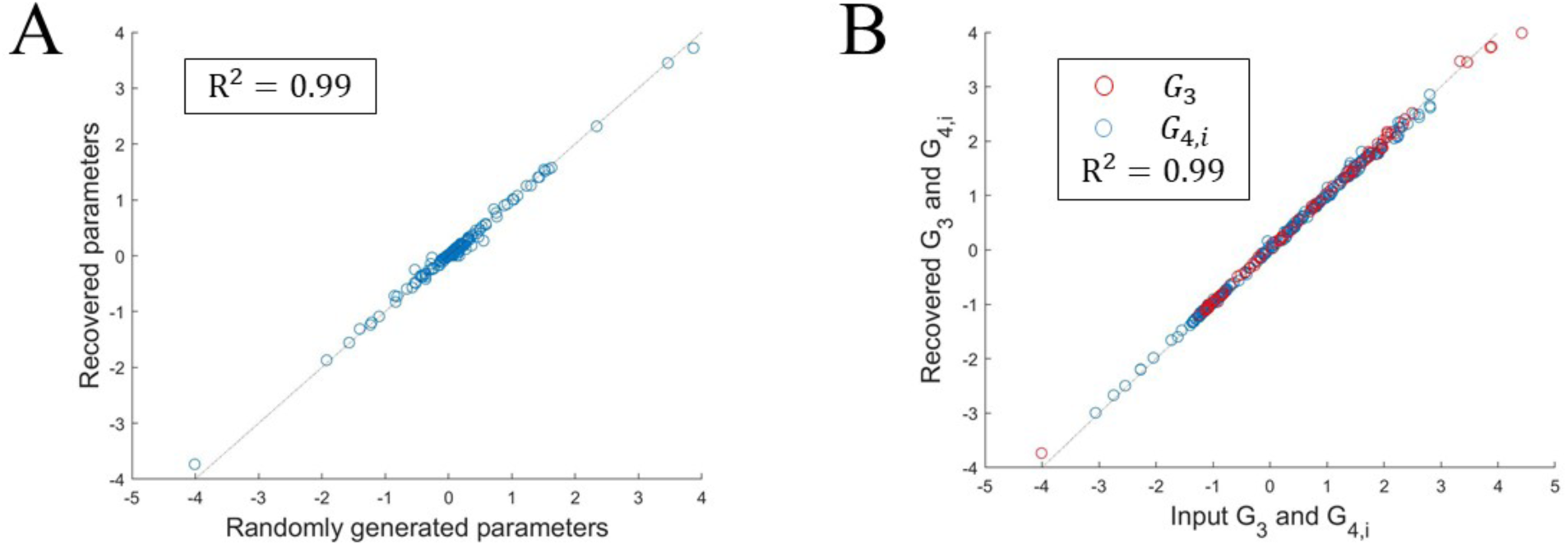
Pairwise model predictions on synthetic data. A set of artificial reaction rates *k*′_*cat,i*_(*S*_*j*_) was generated using a pairwise model with randomly sampled parameters, as described in Materials and Methods. A LASSO fit with cross-validation analogous to that used with real data was employed to recover the parameters of the model. Shown are the comparison between synthetically generated and predicted one- and two-aa model parameters (A) and the corresponding free energy values (B).

**Figure S4.**
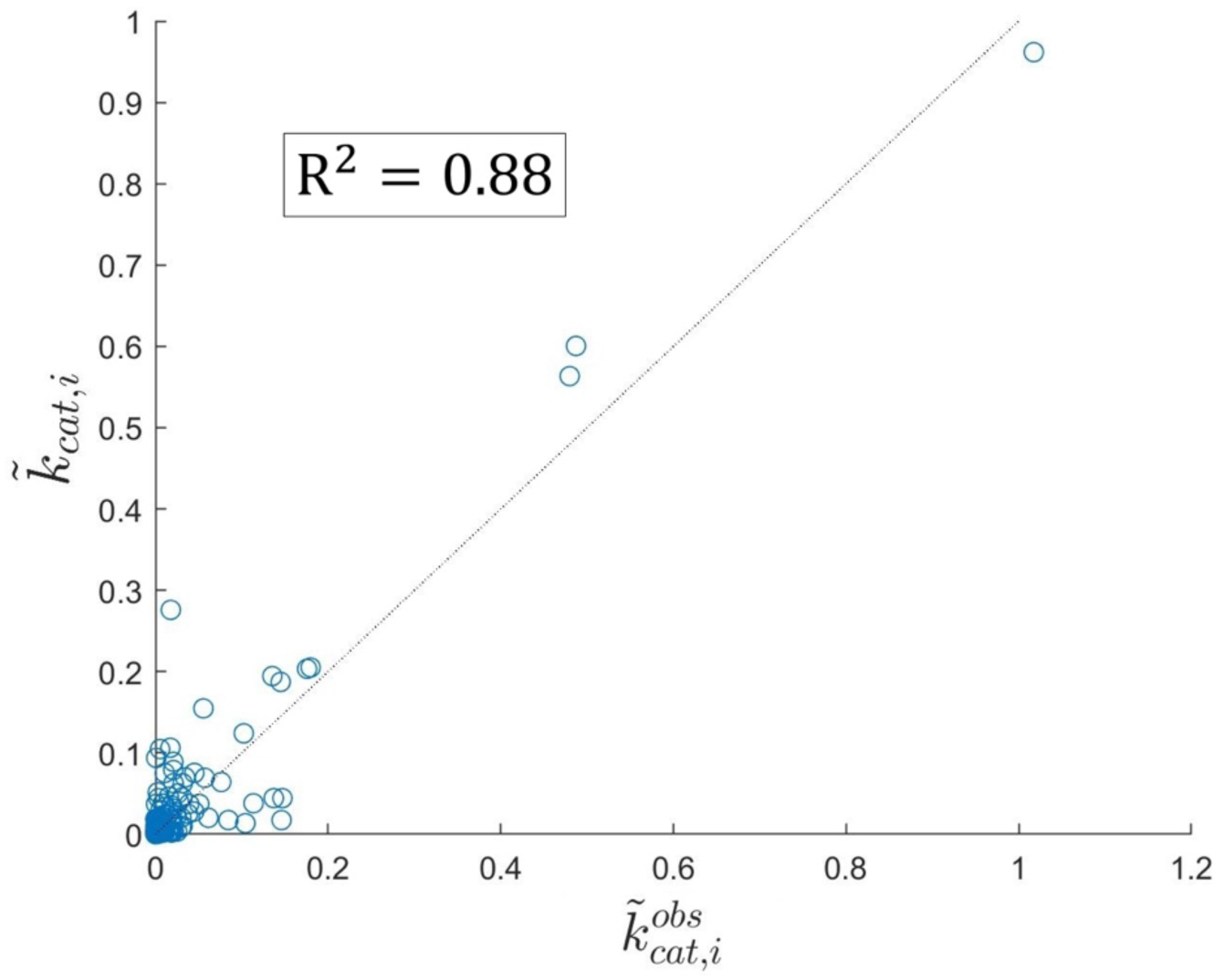
Prediction of novel reaction rates. Comparison of predicted and observed reaction rates *k*_*cat,i*_ for 40 enzyme sequences which were not used in training the model. The parameters of the model were obtained by LASSO with cross-validation using reaction rates for the other 82 enzyme sequences as input data.

**Figure S5.**
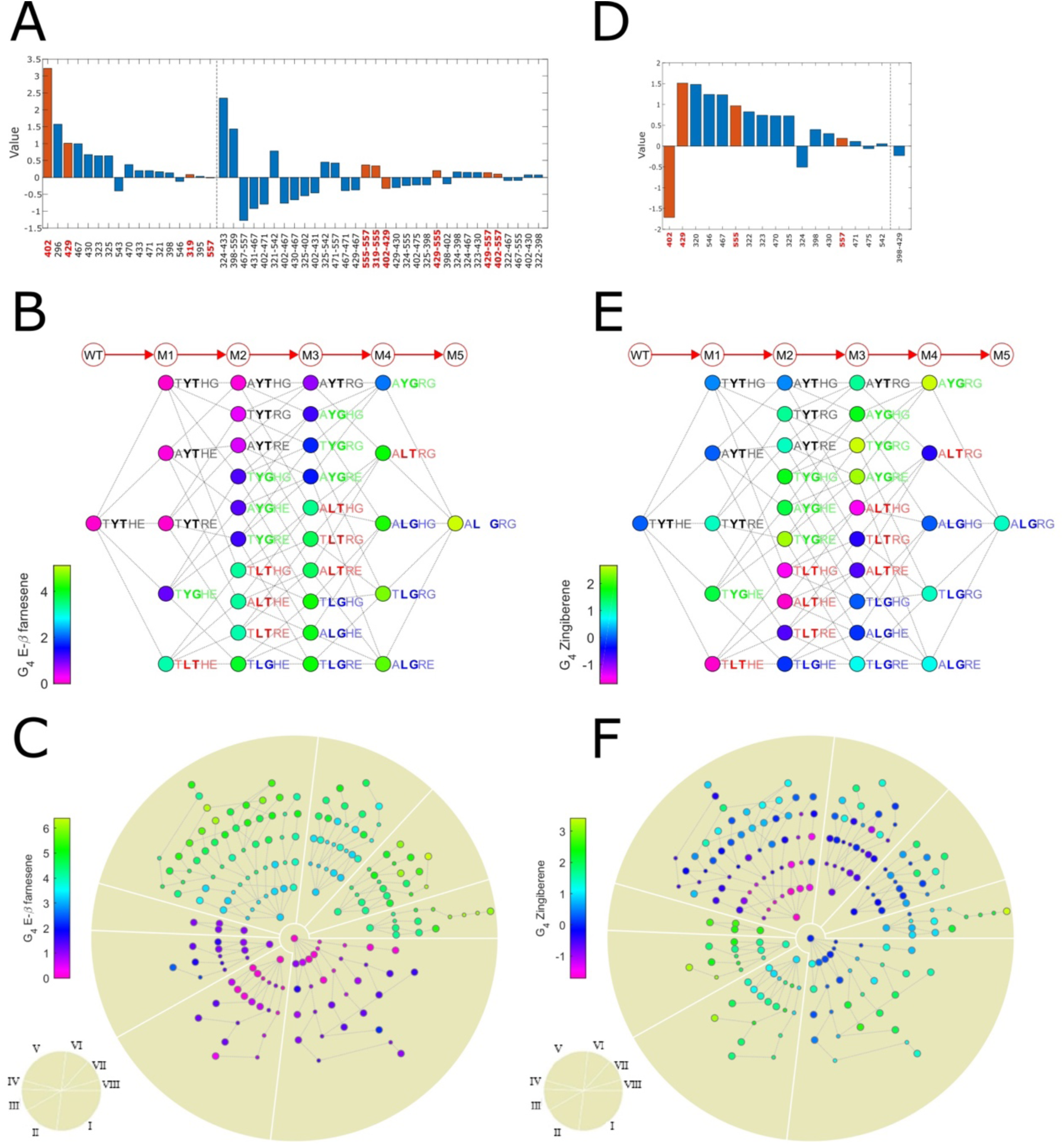
Michaelis-Menten free energy landscapes. Same as Fig. 2 but for E-β-farnesene and zingiberene. All nodes are sorted as in Fig. 3 in order to facilitate visual comparisons. In panels (B) and (E), the nodes are classified into 4 clusters on the basis of amino acids at positions 402 and 429, which have the largest one-aa contributions on the M5 library (red bars in panels (A) and (D)).

**Figure S6.**
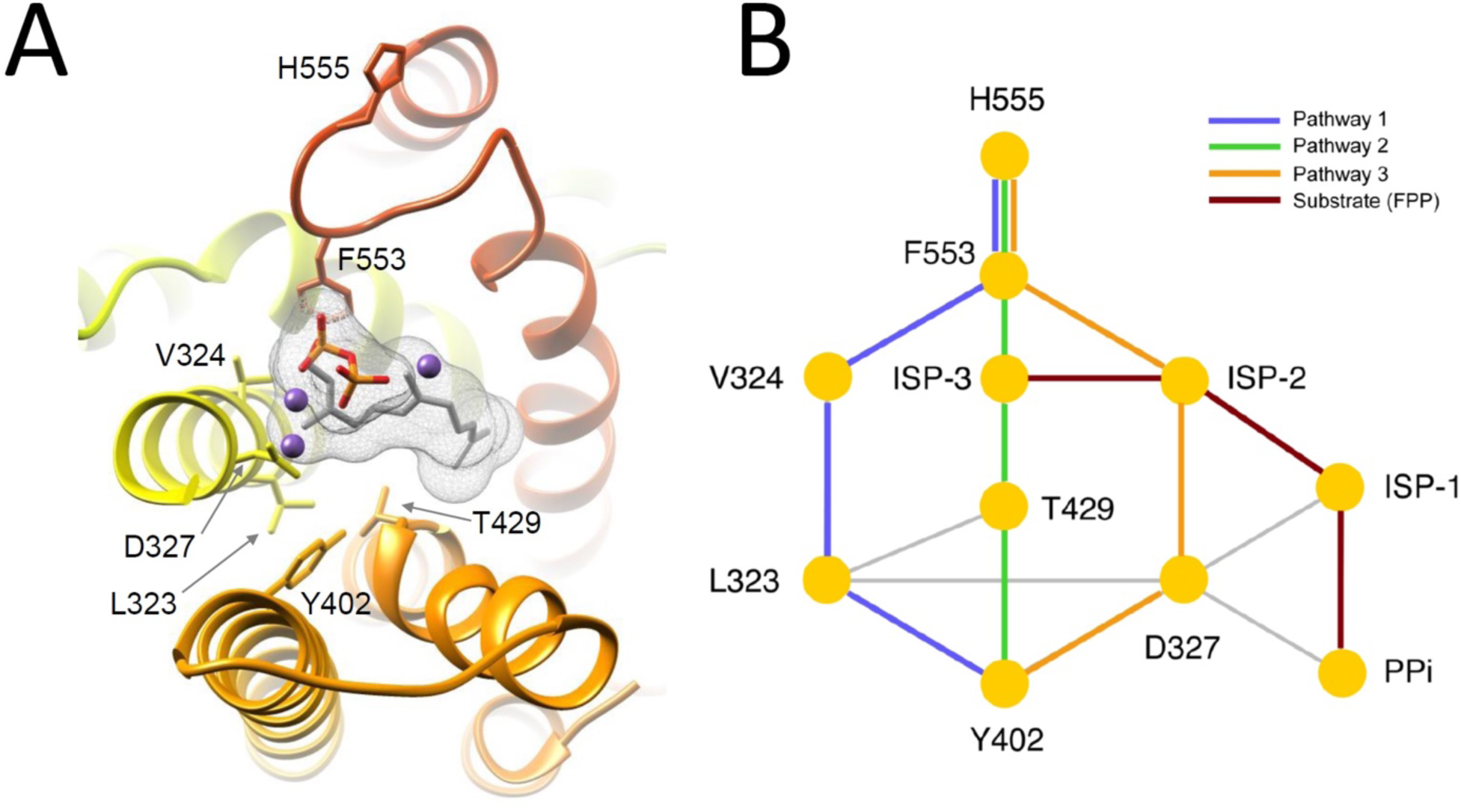
Network analysis of structurally distant interactions in the 402-555 pair. (A) Residues 402 and 555 are highlighted on the BFS structural homology model (created in I-TASSER [55, 56]) with docked FPP substrate (mesh) [1]. Residues involved in interaction networks are labeled. Magnesium ions are shown as purple spheres. (B) Shortest interactions paths between residues 402 and 555 were analyzed in Cytoscape [57]. A network model was constructed from the BFS structural homology model using RINerator [8]. Three shortest interaction paths of identical length were found, as indicated by color. Network edges that are not part of the shortest paths are shown in light grey. Nodes are labeled by aa type and position, by the substrate isoprene unit number (ISP-1,2,3), or as the pyrophosphate moiety (PPi). Contacts with magnesium ions are not shown.

**Figure S7.**
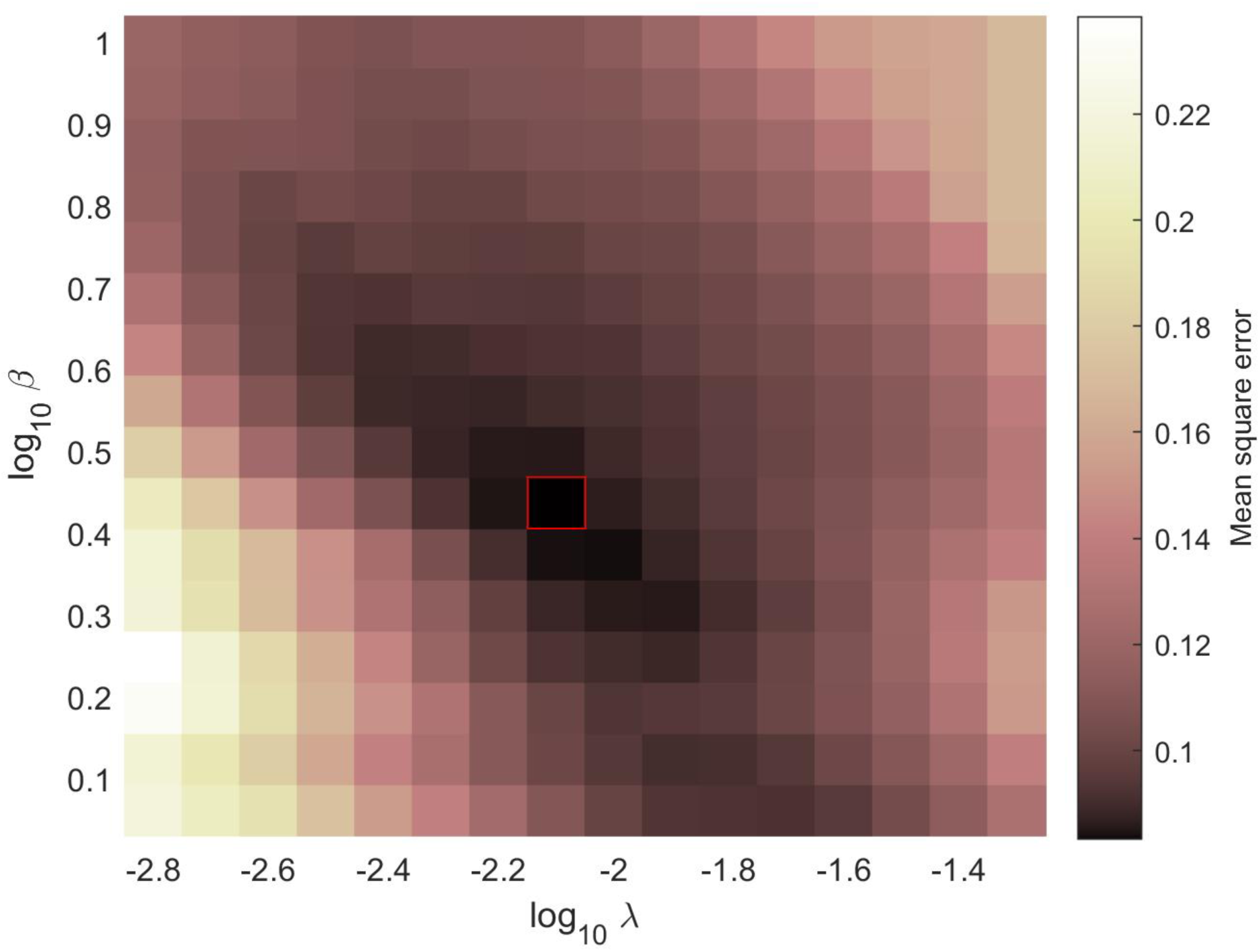
Mean-square error as a function of the LASSO penalty parameters *λ* and *β*. Shown are mean-square errors on the test set obtained via four-fold cross validation (see Materials and Methods for details). The square with the minimum mean-square error, highlighted in red, was used to choose the optimal LASSO parameters.

## Supplementary Table captions

**Table S1. Observed reaction rates 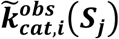 for all enzymes in the combined dataset.** Each row shows the enzyme sequence (*S*_*j*_) at 25 variable positions, followed by 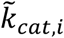 values for 11 products. The first row is the wild-type BFS sequence.

**Table S2. Parameters of the pairwise model fitted on reaction-rate data from the combined library**. (A) One-aa terms for 11 products (*h*_*k*_^(*i*)^) and *G*_3_ (*H*_*k*_) at 25 variable positions. (B) Two-aa terms for 11 products (*ε*_*kl*_^(*i*)^) and *G*_3_ (*E*_*kl*_) for 138 aa pairs. (C) Sequence-independent offsets *C*_*i*_ (Eq. (12)).

**Table S3. Epistatic scores.** (A) Epistatic scores *ES*_*ij*_ for 16 aa pairs, for Michaelis-Menten free energy landscapes and the fitness landscape. Only epistatic scores above the 0.01 threshold are shown. Products omitted from the Table have epistatic scores less than the 0.01 threshold for all aa pairs. Each cell is color-coded to indicate the epistasis type (no epistasis, magnitude epistasis, sign epistasis, reciprocal sign epistasis). Distance refers to *C*_*α*_ − *C*_*α*_ spatial distances (in Å) between amino acids in each pair. (B) Number of aa pairs with no epistasis, magnitude epistasis, sign epistasis, and reciprocal sign epistasis in each free energy and fitness landscape, based on epistatic scores above the 0.01 threshold.

**Table S4. Mass-spec data.** Retention times (RT) and charge-to-mass ratios (m/z) for each product compound considered in this study.

